# Transient contractility attenuation followed by recovery reprograms epithelial cells into a protrusion-driven state that drives tissue fluidization

**DOI:** 10.64898/2026.03.23.713577

**Authors:** Sound WP, Shutong Liu, Nicole Ng, Thanh Phuong Nguyen, Pawan Kumar Mishra, Pratiman De, Aditya Sanjay Gupta, Chantrice Melody Dizon Miranda, Tsuyoshi Hirashima

**Affiliations:** Mechanobiology Institute, National University of Singapore, 5A Engineering Drive 1, Singapore, 117411, Singapore; Department of Physiology, Yong Loo Lin School of Medicine, National University of Singapore, 2 Medical Drive MD9, Singapore, 117593, Singapore

**Keywords:** Collective cell migration, ERK signaling, Mechanobiology, Mechano-chemical feedback, Rigidity transition, Tissue fluidization

## Abstract

Collective cell migration drives tissue morphogenesis, repair and remodeling, and is often accompanied by transitions from solid-like to fluid-like states. While such tissue fluidization has been linked to physical parameters such as cell density, shape and activity, how it is actively regulated by mechano-chemical interplay remains unclear. Previous research has shown that transient attenuation of actomyosin contractility induces a transition from pulsatile, spatially confined motion to coherent, persistent long-range collective flow; however, the underlying cellular and signaling mechanisms remain unclear. Here we uncover the mechanistic basis by which transient perturbation of cell contractility reprograms the migration mode of confluent epithelial cells into a protrusion-driven, fluidizing state, by combining kinase-reporter live imaging, force measurements and mathematical modeling. This transition arises from coordinated changes in cell morphology and mechanics, including reduced cortical tension and greater stretch-induced cell strain, together with enhanced cell–substrate adhesion and force transmission. At the signaling level, this process is accompanied by a rewiring of extracellular signal-regulated kinase (ERK)-mediated mechanotransduction toward a protrusion-coupled mode that sustains migration even under fully confluent conditions. Consistently, a multicellular computational model further demonstrates that protrusion-driven migration is sufficient to promote shape–velocity alignment and drive a transition from caged to flocking-like collective states. Together, our results identify transient contractility attenuation followed by recovery as a trigger for a protrusion-driven state that fluidizes confluent epithelial tissues through coordinated remodeling of cytoskeletal, adhesive, and signaling systems.

## INTRODUCTION

Biological tissues can dynamically transition between distinct physical states, such as solid-like and fluid-like behaviors^1–3^. These transitions arise through coordinated mechanical and topological rearrangements at the cellular level, enabling collective cell movements essential for various biological phenomena. Such state changes, often referred to as tissue fluidization, occur in systems where cells not only generate mechanical forces but also sense and respond to them via biochemical signaling pathways^4–6^. As a result, tissues behave as active, mechano-responsive materials, whose macroscopic properties emerge from tightly coupled mechanical and chemical processes at the cellular scale^7,8^. Physiological manifestations of tissue fluidization include epithelial unjamming during wound healing, large-scale cell rearrangements during morphogenesis, and coordinated collective migration in dynamically remodeling tissues^9–11^. For example, pulsatile actomyosin contractions have been described in developmental epithelia, including Drosophila ventral furrow formation during gastrulation, where contractile pulses drive apical constriction^12^.

The physical basis of tissue fluidization has been extensively explored through theoretical and computational studies, which identify key cellular properties governing collective behavior^13^. Among these, cell density and activity have long been recognized as fundamental parameters, drawing parallels to jamming transitions in non-living systems, where reduced packing or enhanced activity facilitates fluid-like dynamics^14,15^. Building on this general physical framework, subsequent studies have highlighted cell-intrinsic properties unique to living systems. In particular, changes in cell shape, especially elongation, and cell deformability have emerged as critical factors that facilitate neighbor exchanges and promote collective rearrangements^16–20^. These findings suggest that tissue fluidization arises not only from packing conditions, but also from active modulation of cellular geometry and mechanical compliance. However, despite these advances, much of our current understanding remains rooted in theoretical models and analogies to physical systems, and experimental validation in living tissues is still incomplete. In particular, how these physical parameters are dynamically regulated by cell-intrinsic mechanisms, and how they are integrated with intracellular signaling systems, remains largely unresolved.

Experimentally, fluid-like behaviors in epithelial tissues have been observed under various perturbations^21^. In particular, a recent study demonstrated that transient modulation of actomyosin contractility, such as blebbistatin treatment followed by washout, induces a pronounced increase in collective cell motion^22^. The study further showed that epithelial monolayers can switch between distinct collective modes, including pulsatile and flowing states, under such chemical perturbations, which can be captured by minimal coarse-grained frameworks based on polarity alignment and cytoskeletal regulation. Nevertheless, how these perturbations are translated into cell-intrinsic and signaling mechanisms that generate sustained collective flow remains a major gap. Importantly, this response is counterintuitive, as inhibition of myosin II activity is generally expected to suppress traction force generation^23^ and thereby reduce cell migration^24^.

In parallel, a previous study reported that epithelial cells subjected to blebbistatin treatment followed by washout exhibit features reminiscent of leader cells^25^. Together with recent findings that such perturbations can induce large-scale collective flows, these observations raise the possibility that contractility attenuation followed by recovery induces a distinct migratory state sharing selected features with previously described leader cells. Based on this, we hypothesized that blebbistatin washout generates cellular properties resembling those reported for leader cells and whether these changes are associated with collective cell flow. To address this, we combined quantitative experiments and computational simulations, and show that transient attenuation of contractility drives coordinated changes in cell morphology, mechanical properties, and signaling dynamics, thereby establishing a mechanistic link between cell migration mode and tissue -scale fluidization.

## RESULTS AND DISCUSSION

### Blebbistatin washout induces cell enlargement and elongation in epithelial collectives

To test the hypothesis, we established an experimental system based on a previously described protocol^25^. Briefly, Madin–Darby canine kidney (MDCK) cells were treated with blebbistatin for 16 hours, followed by replacement with fresh blebbistatin-free medium for a 4-hour recovery period. Cells were then reseeded at defined densities to initiate subsequent analyses. Cells subjected to this protocol are hereafter referred to as blebbistatin-washout (BWO) cells, whereas cells treated with dimethyl sulfoxide (DMSO) and subjected to the same washout procedure are referred to as DMSO-washout (DWO) cells and used as controls. The time of reseeding was defined as Day 0 (**Figure 1A**).

**Figure 1.**
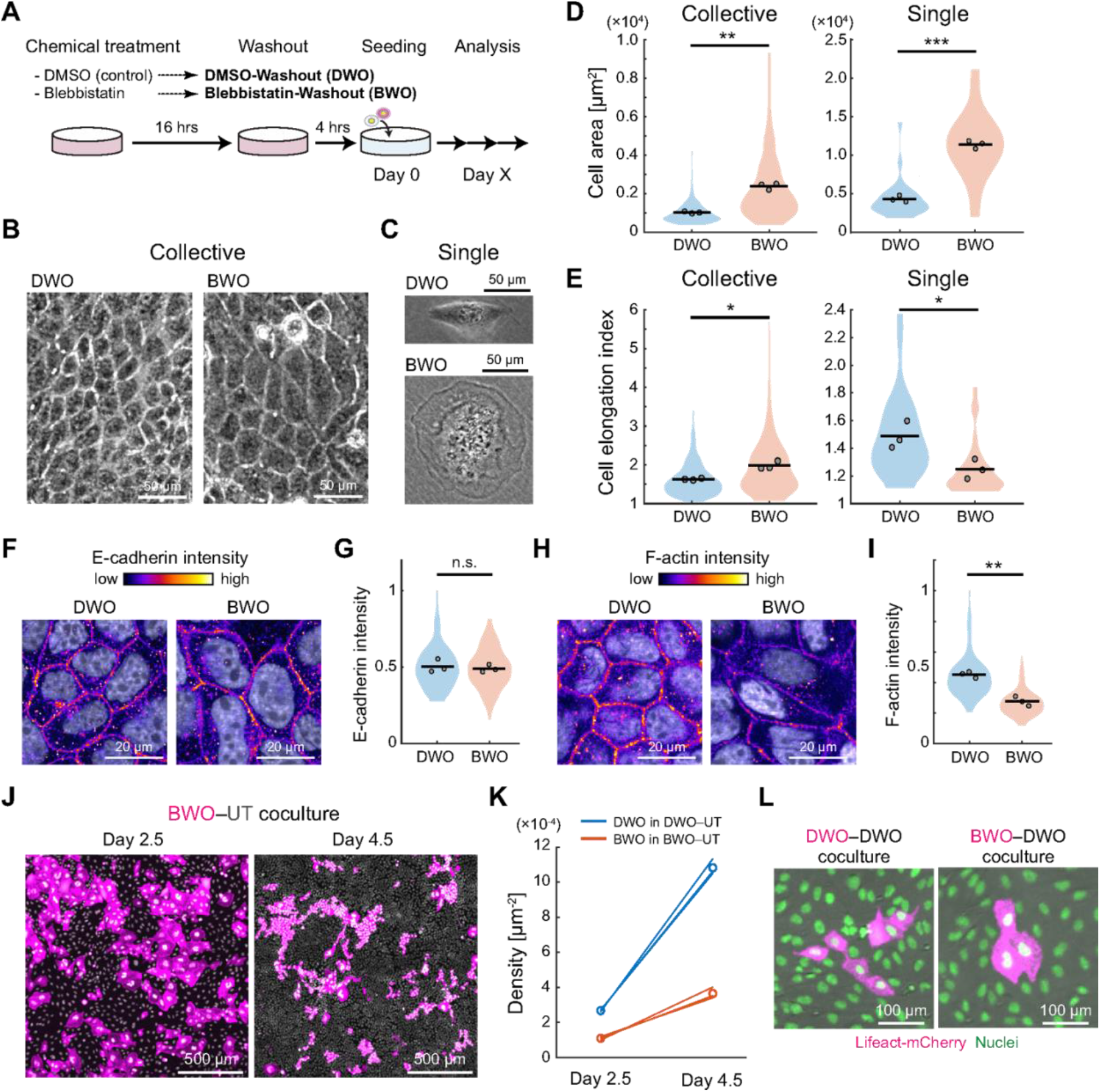
Morphological responses of epithelial cells to blebbistatin washout. (A) Schematic of the experimental workflow. Cells were treated with DMSO (control) or blebbistatin for 16 h, followed by drug washout and a 4-h recovery period to generate DMSO-washout (DWO) and blebbistatin-washout (BWO) cells. The time of seeding was defined as Day 0, and analyses were performed at the indicated time points (Day X). (B,C) Phase-contrast images of DWO and BWO cells in collective monolayers (B) and as isolated single cells (C) on Day 1. Scale bars, 50 µm. (D) Quantification of cell area for DWO and BWO cells in collective (left) and single-cell (right) conditions. Collective: n>340 cells per condition, N=3; Welch’s two-sample t-test on sample means, p=2.79×10^-3^. Single-cell: n=30 cells per condition, N=3; Welch’s two-sample t-test on sample means, p=7.32×10^-5^. (E) Quantification of cell elongation index for DWO and BWO cells in collective (left) and single-cell (right) conditions. Collective: n>340 cells per condition, N=3; Welch’s two-sample t-test on sample means, p=0.0202. Single-cell: n=30 cells per condition, N=3; Welch’s two-sample t-test on sample means, p=0.0353. (F) Immunofluorescence images of E-cadherin with nuclear counterstaining (white) in DWO and BWO cells on Day 1. Scale bars, 20 µm. (G) Normalized membrane E-cadherin intensity, scaled to the maximum value within sample data. n=150 cells per condition, N=3; Welch’s two-sample t-test on sample means, p=0.641. (H) Phalloidin staining showing F-actin distribution with nuclear counterstaining (white) in DWO and BWO cells on Day 1. Scale bars, 20 µm. (I) Normalized cortical actin intensity, scaled to the maximum value within sample data. n=150 cells per condition, N=3; Welch’s two-sample t-test on sample means, p=2.36×10^-3^. (J) Representative images for the co-culture assay of BWO and untreated (UT) cells at Day 2.5 (left) and at Day 4.5 (right). BWO cells were labeled with Lifeact-mCherry. Scale bars, 500 µm. (K) Change in nuclear density of DWO (blue) and BWO (red) cells in co-culture from Day 2.5 to Day 4.5. Lines represent individual samples and open circles indicate mean values at each timepoint. N=3. (L) Fluorescence images of homotypic DWO-DWO co-culture (left) and heterotypic BWO-DWO co-culture (right) at Day 2.5. In the homotypic condition, one population was labeled with Lifeact-mCherry; in the heterotypic condition, BWO cells were labeled. Scale bars, 100 µm.

BWO treatment induced pronounced and context-dependent morphological changes by Day 1. Under confluent conditions, BWO cells displayed a markedly expanded and elongated morphology compared with DWO cells (**Figure 1B**). In contrast, when examined as isolated single cells, BWO cells adopted a more rounded morphology than DWO cells (**Figure 1C**). Quantitative analysis revealed that BWO cells exhibited an approximately 146% increase in cell area relative to DWO cells under confluent conditions, and a 164% increase under isolated single-cell conditions (**Figure 1D**). This enlargement was spatially heterogeneous, with expanded cells coexisting with less-expanded neighboring cells, rather than reflecting uniform expansion of all cells across the monolayer. Notably, the cell elongation, defined as the inverse of circularity, was increased by 20% in BWO cells compared with DWO cells in confluent monolayers, whereas it was reduced by 16% in isolated BWO cells relative to isolated DWO cells (**Figure 1E**). These results indicate that BWO treatment enhances autonomous cell spreading on the substrate, while cell elongation is preferentially promoted only in the presence of cell–cell interactions.

Since epithelial cell morphology is governed by the balance between intercellular mechanical coupling mediated through adhesions and intracellular cortical tension^16,26,27^, we next examined the distribution of the main cell-cell adhesion and cytoskeletal components, E-cadherin and F-actin on Day 1. Fluorescence staining revealed no significant difference in E-cadherin localization or intensity between DWO and BWO cells (**Figures 1F,G**), indicating that adherens junction integrity is largely preserved following BWO treatment. In contrast, cortical actin at the lateral side of cells was markedly reduced in BWO cells compared with DWO controls (**Figures 1H,I**). These observations suggest that BWO treatment weakens cortical tension without substantially altering cell–cell adhesion strength. In this configuration, reduced cortical actomyosin tension would lower the resistance to cell-shape deformation, while preserved cell–cell adhesion would maintain mechanical coupling between neighboring cells, allowing anisotropic intercellular forces to elongate cells within the monolayer. Such a mechanical configuration is consistent with theoretical predictions of elongated epithelial morphology^16,17,28^.

Based on previous work showing that leader cells can be eliminated from epithelial populations by mechanical cell competition^25^, we asked whether BWO cells would show a similar competitive disadvantage when co-cultured with untreated (UT) control cells. Characterization of the BWO population, however, indicated only partial correspondence with the previously described p53/p21-associated leader-cell population: BWO cells showed only a small multinucleated fraction (∼5%) and no increase in phospho-p53 nuclear enrichment, while p21 levels largely overlapped with DWO controls, despite a subset of cells exhibiting elevated nuclear p21 (**Figure S1**). Consistent with this distinction, BWO cells were not selectively eliminated by UT cells.

Instead, they persisted over extended culture periods and remained stably integrated within the epithelial monolayer (**Figure 1J**). The number of BWO cells increased by 3.34-fold over two days in BWO–UT co-cultures, although the rate of increase was lower than that observed for DWO cells in DWO–UT co-cultures (**Figure 1K**). Moreover, BWO cells surrounded by DWO cells in heterotypic co-culture exhibited a more expanded morphology compared with DWO cells in homotypic DWO–DWO co-cultures (**Figure 1L**). This observation suggests that BWO cells generate stronger cell-spreading forces than DWO cells during heterotypic cell–cell interactions, rather than being mechanically disadvantaged in the space-competitive epithelial environment.

### BWO enhances cell–substrate adhesion and force transmission

Since BWO cells exhibited enhanced cell spreading (**Figures 1D,L**) and resistance to mechanical cell competition (**Figures 1J,K**), we next asked whether BWO enhances cell–substrate adhesion. As an initial and simple functional readout, we quantified cell detachment kinetics in response to trypsin treatment (**Figure 2A**). Time-lapse imaging revealed that, upon trypsinization, approximately 50% of DWO cells detached from the substrate within 10 minutes, whereas only 6% of BWO cells were detached at the same time point (**Figure 2B**). This marked delay in detachment suggests that BWO cells adhere more strongly to the underlying substrate than control cells. This difference in trypsin-detachment kinetics was no longer detectable by Day 2 and remained absent at later time points (**Figure S2**), indicating that this adhesion-related BWO phenotype is reversible rather than permanently maintained. To directly quantify adhesive strength, we employed a centrifugation-based adhesion assay, in which a defined centrifugal force is applied to detach adherent cells. Because MDCK monolayers remained attached even at the maximum centrifugal force available in our setup, this assay was performed using HEK293 cells as an experimentally tractable readout of BWO-dependent changes in cell–substrate adhesion. Under this regime, the force required to detach cells provides a direct measure of cell–substrate adhesive force^29–31^. Cells were plated in parallel and allowed to adhere, after which each sample was subjected to a single defined centrifugal force for 3 min. In both DWO and BWO conditions, the resulting curves exhibited characteristic saturating profiles, with an approximately one-order shift in the force range **(Figure 2C**). The centrifugal force required to detach 50% of cells was 15-fold higher for BWO cells than for DWO controls (**Figure 2D**).

**Figure 2.**
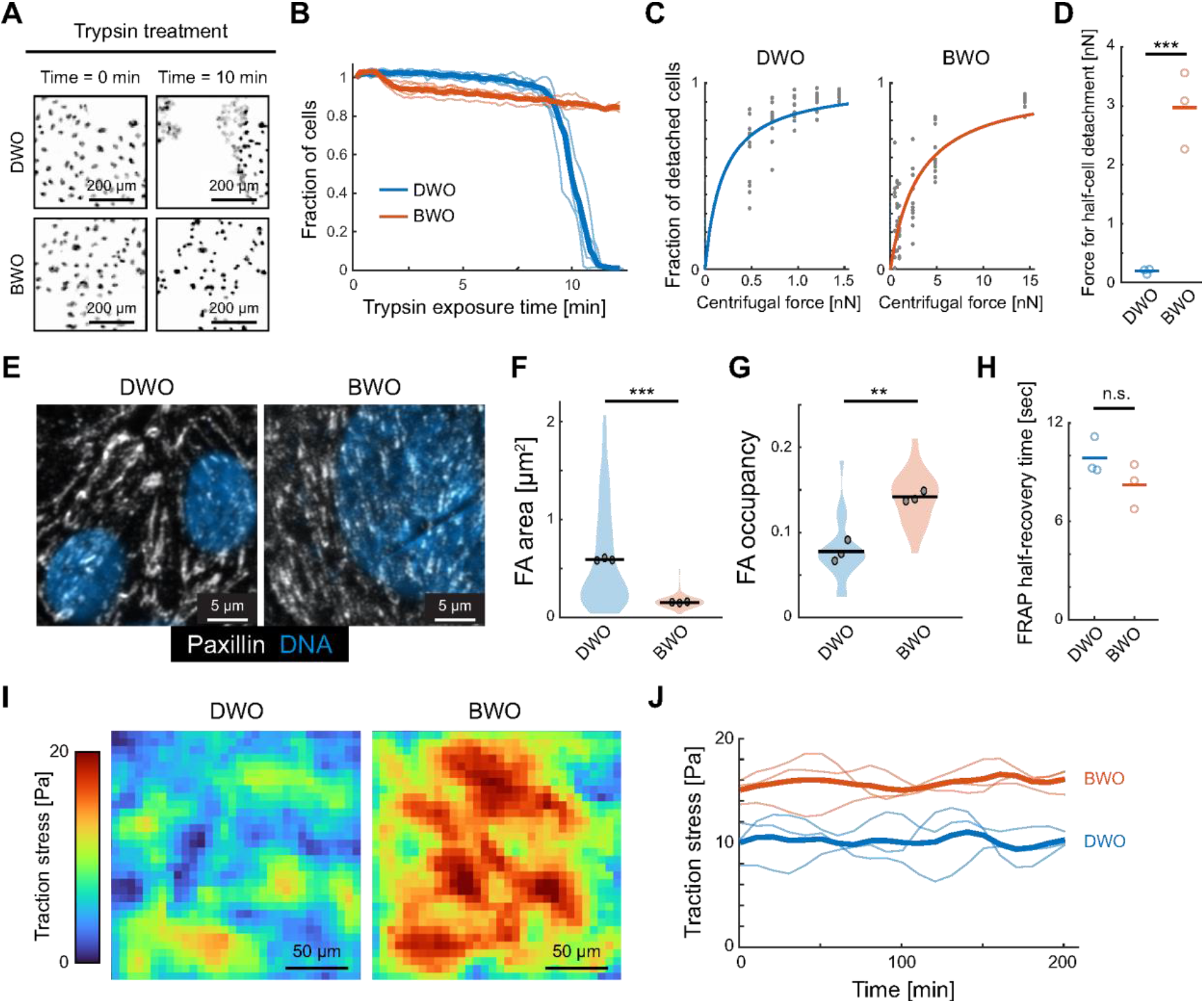
Measurement of cell–substrate adhesion. (A) Representative still images of DWO and BWO cells at the onset of trypsin treatment (left) and after 10 min of exposure (right). Nuclei are shown in gray. Scale bars, 200 µm. (B) Fraction of cells remaining attached to the substrate during trypsin treatment for DWO (blue) and BWO (red) conditions as a function of exposure time. Thick lines indicate mean values. N=4. (C) Fraction of detached HEK293 cells measured by the centrifugation assay for DWO (left) and BWO (right) cells as a function of applied centrifugal force. Solid lines represent fitted detachment curves. n≥12 per condition; N=3. (D) Centrifugal force required for half-maximal detachment of HEK293 cells for DWO and BWO conditions. N=3; two-sample t-test, p=1.80×10^-3^. (E) Immunofluorescence images of paxillin (white) with nuclear counterstaining (blue). Scale bars, 5 µm. (F,G) Quantification of focal adhesion (FA) properties, including FA area (F) and FA occupancy within individual cells (G). FA area: n≥196 per condition, N=3; Welch’s two-sample t-test on replicate means, p=8.04×10^-4^; FA occupancy: n=30 per condition, N=3; Welch’s two-sample t-test on replicate means, p=4.41×10^-3^. (H) FRAP half-recovery time of paxillin in DWO and BWO cells. N=3; two-sample t-test, p=0.191. (I) Spatial maps of traction stress magnitude for DWO (left) and BWO (right) cell collectives at Day 1.5. Scale bars, 50 µm. (J) Mean traction stress of DWO (blue) and BWO cells (red) as a function of time. Thick lines indicate mean values. N=3.

We next examined whether the enhanced cell–substrate adhesion observed in BWO cells was accompanied by changes in focal adhesion (FA) organization. Paxillin was used as canonical markers of FAs^32,33^. Immunofluorescence analysis revealed clear differences in FA architecture between the two conditions (**Figure 2E**). Specifically, BWO cells displayed significantly smaller individual FAs with reduced size variance compared with DWO cells (**Figure 2F**). In contrast, the overall FA occupancy within individual cells, defined as the total FA area normalized to cell area, was significantly increased in BWO cells relative to controls (**Figure 2G**). Individual FA size is not necessarily a direct proxy for global cell–substrate adhesion strength, which depends on the overall organization and extent of the adhesive interface^34^. Increased cell spreading can also be accompanied by an increase in total FA area despite a decrease in mean individual FA area^35^. Thus, the smaller individual FAs together with increased FA occupancy in BWO cells indicate a redistribution of FA architecture toward greater overall adhesive coverage. Together with the independent detachment measurements, these findings suggest that enhanced adhesion in BWO cells is associated with altered organization of the cell–substrate adhesive interface rather than enlargement of individual FAs. Further, to assess whether BWO alters FA molecular dynamics, we performed fluorescence recovery after photobleaching (FRAP) experiments on paxillin to quantify FA protein turnover. Analysis of FRAP recovery curves revealed no significant difference in the half-time of fluorescence recovery between DWO and BWO cells (**Figure 2H**), indicating that paxillin binding dynamics at FAs are largely preserved following BWO treatment.

Finally, we measured cellular traction stress under confluent conditions to assess changes in force transmission to the substrate in BWO cells. Consistent with their increased adhesion strength, BWO cells exhibited an average increase of 15% in traction stress compared with DWO cells (**Figures 2I,J**). These results indicate that BWO enhances the ability of epithelial cells to transmit mechanical forces to the underlying substrate within a collective context. Notably, cellular traction stresses were measured in confluent monolayers, where force generation reflects not only cell–substrate interactions but also mechanical coupling between neighboring cells. Thus, the increased traction observed in BWO cells likely arises from a combination of enhanced cell –substrate adhesion and altered force balance within the epithelial sheet, rather than from autonomous changes in single-cell contractility alone. Accordingly, the large shift in detachment force and the more modest increase in traction stress should not be interpreted as directly proportional quantities, because they reflect distinct mechanical readouts measured in different cellular contexts: adhesive failure of HEK293 cells under externally imposed loading versus active force transmission within confluent MDCK monolayers.

### BWO accelerates monolayer expansion and increases stretch-induced cell strain

The morphological and mechanical differences observed in BWO cells prompted us to investigate how BWO influences collective epithelial migration. To this end, we performed a monolayer expansion assay, in which a pre-confined epithelial sheet is released to expand into free space. Upon release of confinement at Day 1.5, DWO monolayers migrated into the free space with a characteristic spatiotemporal pattern as anticipated. Specifically, cells at the leading edge advanced forward while multiple velocity waves propagated from the front toward the bulk of the tissue (**Figures 3A,B, top; Video 1**). These velocity waves were accompanied by pulsatile movements of bulk cells, consistent with previously described collective migration dynamics^36,37^. In contrast, BWO monolayers displayed a markedly different behavior; an initial velocity propagation rapidly traversed the tissue following confinement release, after which the monolayer continued to migrate at a relatively constant speed without pronounced oscillatory behavior (**Figures 3A,B, bottom; Video 1**). Notably, the leading-edge speed of BWO monolayers was approximately 35% higher than that of DWO monolayers (**Figure 3C**).

**Figure 3.**
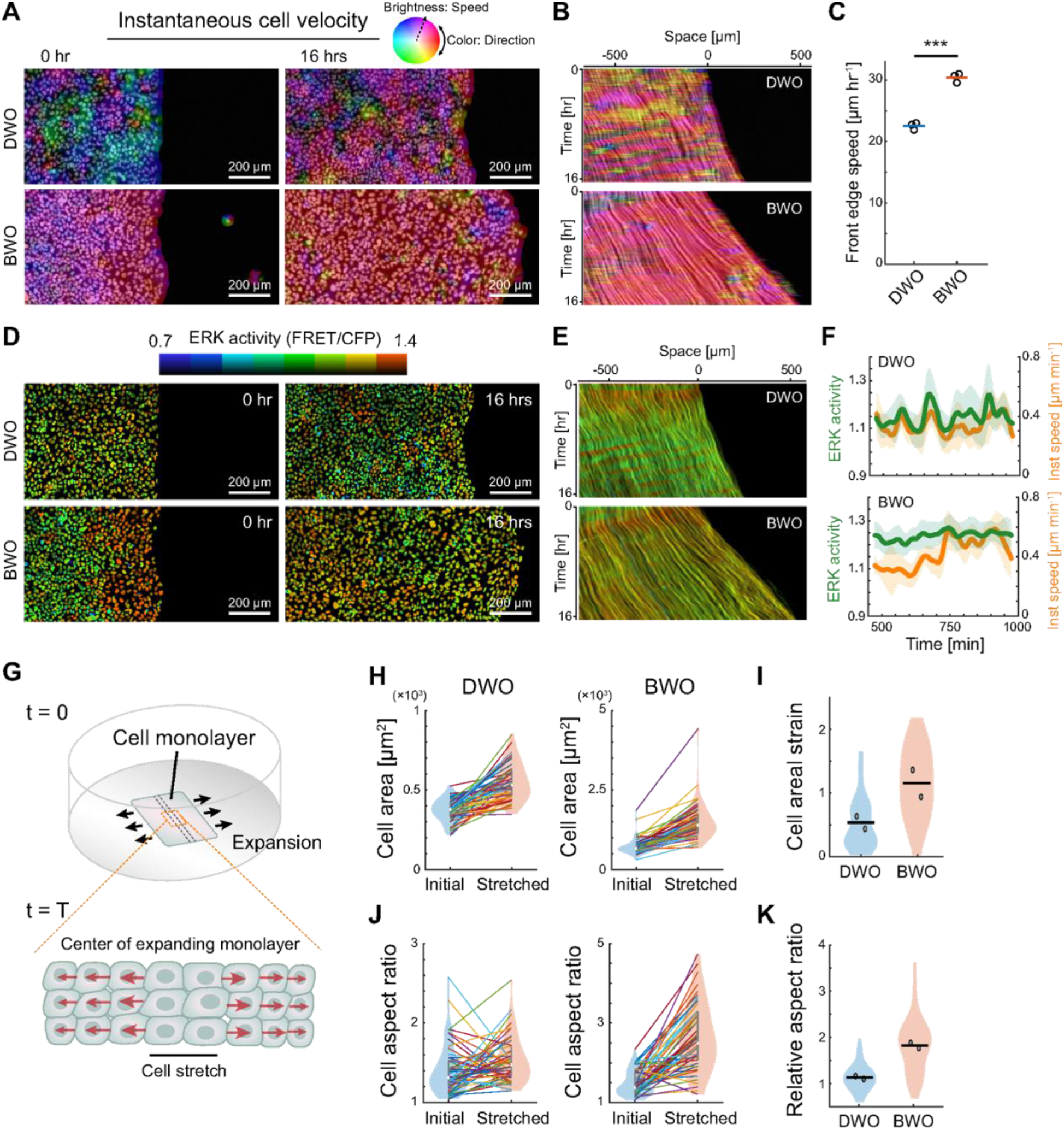
BWO-induced tissue expansion and stretch-induced cell strain. (A) Representative still images of cell velocity fields (rainbow color) superimposed on nuclear images (gray) for DWO (top) and BWO (bottom) monolayers at the onset of expansion (left) and 16 h after release (right). Color indicates velocity direction, and brightness represents speed. Scale bars, 200 µm. (B) Kymographs of cell velocity for DWO (top) and BWO (bottom) monolayers, displayed as velocity maps (rainbow color) overlaid on nuclear images. The spatial origin on the horizontal axis corresponds to the initial position of the leading edge. (C) Leading-edge speed of DWO and BWO monolayers, quantified over the time window from 2 to 14 h after release. N=3; two-sample t-test, p=2.14×10^-4^. (D) Representative still images of ERK activity maps for DWO (top) and BWO (bottom) monolayers at the onset of expansion (left) and 16 h after release (right), corresponding to panel (A). Scale bars, 200 µm. (E) Kymographs of ERK activity for DWO (top) and BWO (bottom) monolayers, corresponding to panel (B). The spatial origin on the horizontal axis indicates the initial position of the leading edge. (F) ERK activity (green) and instantaneous cell speed (orange) at the single-cell level in the bulk region of expanding DWO (top) and BWO (bottom) monolayers. Thick lines denote mean values and shaded regions indicate standard deviation. n>30 cells for both conditions. (G) Schematics of spontaneous cell-stretch assay during monolayer expansion. (H) Cell area measured at the onset of expansion and at the time of maximal cell stretching for DWO (left) and BWO (right) cells. n=50 cells per condition, N=2. (I) Maximum cell area strain in DWO and BWO cells during cell-stretch assay. n=50 cells per condition, N=2. (J) Cell aspect ratio measured at the onset of expansion and at the time of maximal cell stretching for DWO (left) and BWO (right) cells. n=50 cells per condition, N=2. (K) Relative change in cell aspect ratio for DWO and BWO cells. n=50 per condition, N=2.

Given that collective epithelial migration is tightly coupled to extracellular signal–regulated kinase (ERK) activity^38–40^, we next examined whether the distinct velocity patterns observed in DWO and BWO monolayers were reflected in corresponding ERK activity dynamics. To this end, we performed time-lapse imaging using a Förster resonance energy transfer (FRET)-based ERK biosensor expressed in the cells^41,42^. In DWO monolayers, ERK activity exhibited multiple propagating waves that were temporally and spatially correlated with velocity waves ( **Figures 3D–F, top; Video 2**), consistent with ERK-mediated mechanochemical feedback mechanisms underlying collective migration^43,44^. In contrast, BWO monolayers displayed a single, initial ERK activity propagation following confinement release, followed by a relatively uniform and sustained ERK activation throughout the bulk of the tissue during expansion (**Figures 3D–F, bottom; Video 2**). Given that ERK-driven actomyosin contractility is a key component of the feedback loop that generates oscillatory velocity and ERK waves^45^, the absence of repeated ERK oscillations in BWO monolayers suggests a functional decoupling between ERK activation and contractile force generation. These results indicate that stretch-induced ERK activation remains intact, whereas ERK-dependent contractile responses are attenuated^45,46^. Consequently, although mechanical pulling forces from leading cells are transmitted to follower cells during expansion^47,48^, the resulting contractile response appears insufficient to suppress ERK activity, leading to sustained ERK activation and smoother collective migration dynamics.

The monolayer expansion assay further suggested that BWO cells may undergo greater strain in response to intercellular pulling forces, consistent with the reduced cortical actin observed in BWO cells (**Figure 1I**). To directly test this possibility, we analyzed cell shape changes within the central region of the expanding monolayer, where tensile forces accumulate symmetrically from both sides (**Figure 3G**). For each cell, we identified the time point at which maximal stretch occurred during monolayer expansion and quantified changes in cell area and cell elongation index relative to the initial state. The timing of maximal stretch did not differ significantly between DWO and BWO cells (**Figure S3**). DWO cells exhibited a median increase in cell area of 34%, whereas BWO cells showed a markedly larger median increase of 118% (**Figures 3H,I**). Similarly, the median cell elongation index increased by 12% in DWO cells but by 87% in BWO cells (**Figures 3J,K**). These results demonstrate that BWO cells undergo greater area and shape strain along the axis of spontaneously generated stretch during collective migration. Because BWO cells also differ in their initial size and potentially in their packing state, these measurements should be interpreted as an emergent response of the BWO monolayer rather than as an isolated measure of intrinsic cell deformability.

### BWO induces coherent collective motion and enhanced topological rearrangement in confluent monolayers

We further investigated collective cell motion under confluent conditions by analyzing supracellular-scale dynamics together with single-cell tracking. In DWO monolayers, cells exhibited relatively solid-like behavior: individual cells fluctuated around fixed positions with limited net displacement, consistent with a jammed epithelial state. These local cell movements were accompanied by ERK activity dynamics but did not give rise to coherent long-range motion (**Figures 4A,B, top; Videos 3,4**). In contrast, BWO monolayers displayed markedly more coherent and persistent collective motion. Single-cell trajectories formed coherent paths at supracellular scales aligned with the elongated cell axis and persisting for several hours, indicative of sustained, coherent collective flow (**Figures 4A,B, bottom; Videos 3,4**). The instantaneous cell speed associated with this collective motion was dynamically coupled to ERK activity, with single-cell cross-correlation analysis showing that BWO cells retained ERK–speed coupling comparable to that observed in DWO cells (**Figure S4A,B),** suggesting that ERK signaling participates in coordinating active collective flow in BWO cells. BWO cells exhibited an 89% increase in instantaneous speed relative to DWO cells (**Figure 4C**), consistent with increased traction stress under confluent conditions **(Figures 2I,J)**.

**Figure 4.**
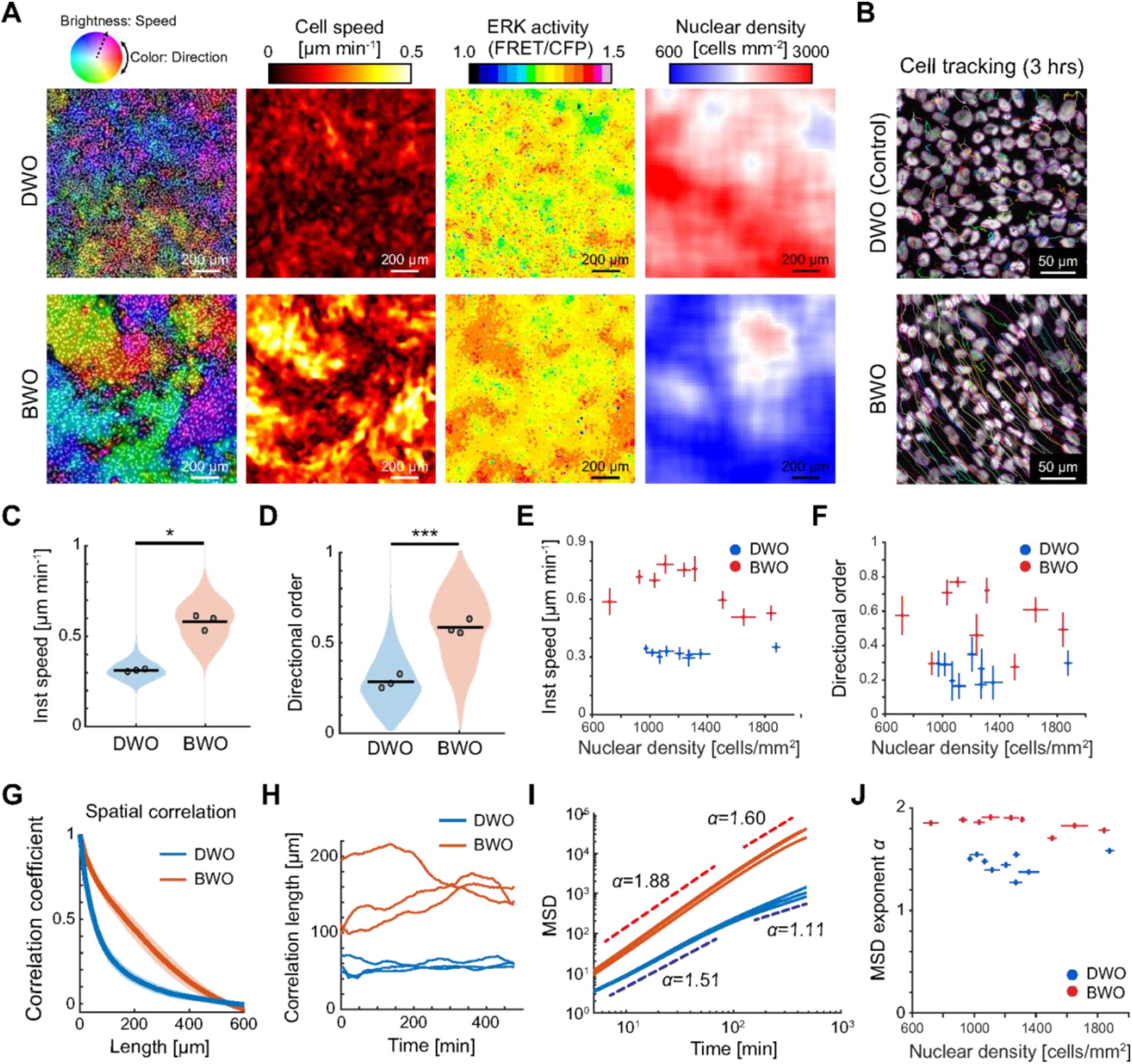
BWO-induced tissue fluidity in monolayer. (A) Representative still images of cell velocity vector fields (rainbow color) overlaid on nuclear images (gray; left), cell speed maps (middle), and ERK activity maps (right) for DWO (top) and BWO (bottom) monolayers. Scale bars, 200 µm. (B) Trajectories of individual cells tracked over 3 h in confluent DWO (upper) and BWO (bottom) monolayers. Cell images correspond to the initial time point. Colored lines indicate trajectories originating from distinct nuclear positions. Scale bars, 50 µm. (C) Time-averaged instantaneous cell speed over an 8-h interval for DWO and BWO monolayer. DWO: n=3421 cells, N=3; BWO: n=3891 cells, N=3; Welch’s two-sample t-test on biological-replicate means, p= 7.38×10^-3^. (D) Directional order parameters quantifying the degree of alignment in collective cell motion for DWO and BWO monolayers. DWO: n=3421 cells, N=3; BWO: n=3891 cells, N=3; Welch’s two-sample t-test on biological-replicate means, p=7.69×10^-4^. (E,F) Density-matched analysis of time-averaged instantaneous cell speed (E) and directional order (F) as a function of nuclear density in local regions. Each point represents one local field of view; error bars indicate s.d. over the 150-min observation period. DWO: n=9 regions, N=3; BWO: n=9 regions, N=3. (G) Representative spatial velocity correlation functions for DWO and BWO monolayers. Thick lines denote mean values, and shaded regions represent standard deviation (s.d.) across all cells at a given time point. (H) Spatial velocity correlation length as a function of time for DWO and BWO monolayers. N=3. (I) Mean squared displacement (MSD) averaged over single-cell trajectories as a function of time. Solid lines indicate experimental measurements, and dotted lines represent power-law fits used to extract scaling exponents. DWO: n=3421 cells, N=3; BWO: n=3891 cells, N=3. (J) MSD scaling exponent α as a function of nuclear density for the same local regions analyzed in (E,F). Each point represents one local field of view. DWO: n=9 regions, N=3; BWO: n=9 regions, N=3.

To assess directional coordination within the tissue, we calculated the directional order parameter, which quantifies the degree of alignment in cell motion. BWO monolayers exhibited significantly higher directional order than DWO monolayers (**Figure 4D**), indicating enhanced coherence of collective movement. Because BWO monolayers also exhibited a lower average nuclear density and greater spatial heterogeneity in cell density than DWO monolayers (**Figures S4C,D**), we next asked whether these differences in collective dynamics could be explained by cell density alone. We therefore analyzed local regions spanning overlapping nuclear-density ranges. Across this overlapping range, BWO cells maintained higher instantaneous speed and directional order than DWO cells (**Figures 4E,F**), indicating that these differences cannot be attributed simply to lower cell density in BWO monolayers.

We next quantified the spatial extent of coordinated motion by measuring the spatial velocity correlation and its correlation length, which reports the characteristic length scale over which cell movements are correlated. BWO monolayers exhibited more slowly decaying spatial velocity correlations than DWO monolayers (**Figure 4G**), corresponding to a consistently longer correlation length across the observation period (**Figure 4H**) and indicating enhanced long-range coordination across the epithelial sheet. Notably, the correlation length in BWO monolayers also showed greater temporal variability than in DWO monolayers (**Figure 4H**). This increased variance suggests that BWO cells reside in a dynamically evolving collective state, in which coherent motion domains continuously emerge, reorganize, and dissipate over time.

Furthermore, we calculated the mean squared displacement (MSD) to assess the persistence of single-cell motion within the collective context (**Figure 4I**). BWO cells exhibited significantly higher motility persistence than DWO cells across both short (<100 min) and long (>100 min) timescales. Consistent with the density-matched speed and directional-order analyses, the MSD scaling exponent remained higher in BWO cells across overlapping nuclear-density ranges (**Figure 4J**). Specifically, BWO cells maintained super-diffusive behavior (α = 1.6–1.9) even at long times, whereas DWO cells exhibited only transient persistence at short times (α = 1.51) that decayed toward near-diffusive behavior at longer times (α = 1.11), consistent with the oscillatory period of ERK activity reported previously^43,44^. These results indicate that BWO stabilizes persistent, directed motion within confluent epithelial collectives. BWO cells also exhibited greater neighbor loss over 120 min and a higher rate of persistent T1-like Delaunay edge flips than DWO cells (**Figures S4E–F**), providing direct evidence for enhanced topological rearrangement in addition to coherent collective motion.

### BWO rewires ERK signaling toward an HGFR-dependent mode

The morphological and mechanical features observed in BWO cells, including enlarged and elongated morphology, elevated traction forces, and enhanced migratory activity, overlap with features reported for epithelial leader cells^24,49–51^. We therefore asked whether BWO treatment also alters the upstream signaling architecture controlling ERK activity toward a pattern resembling that reported in leader cells. Previous work has demonstrated that ERK signaling in leader cells is predominantly regulated by hepatocyte growth factor receptor (HGFR/c-Met), whereas non-leader or follower cells rely primarily on epidermal growth factor receptor (EGFR)–dependent regulation^52^. Under fully confluent conditions, where no free space is available for leader cells to emerge, we tested whether BWO cells exhibit HGFR-dependent ERK signaling.

As an initial assessment, we examined HGFR activation by immunostaining for phosphorylated HGFR at its catalytic domain. Activated HGFR was detected at the plasma membrane in both cell types, but its signal intensity was markedly higher in BWO cells than in DWO controls (**Figure 5A**). Quantification revealed that membrane-associated phosphorylated HGFR levels were increased by 1.48-fold in BWO cells relative to DWO cells (**Figure 5B**), suggesting BWO enhanced HGFR signaling activity.

**Figure 5.**
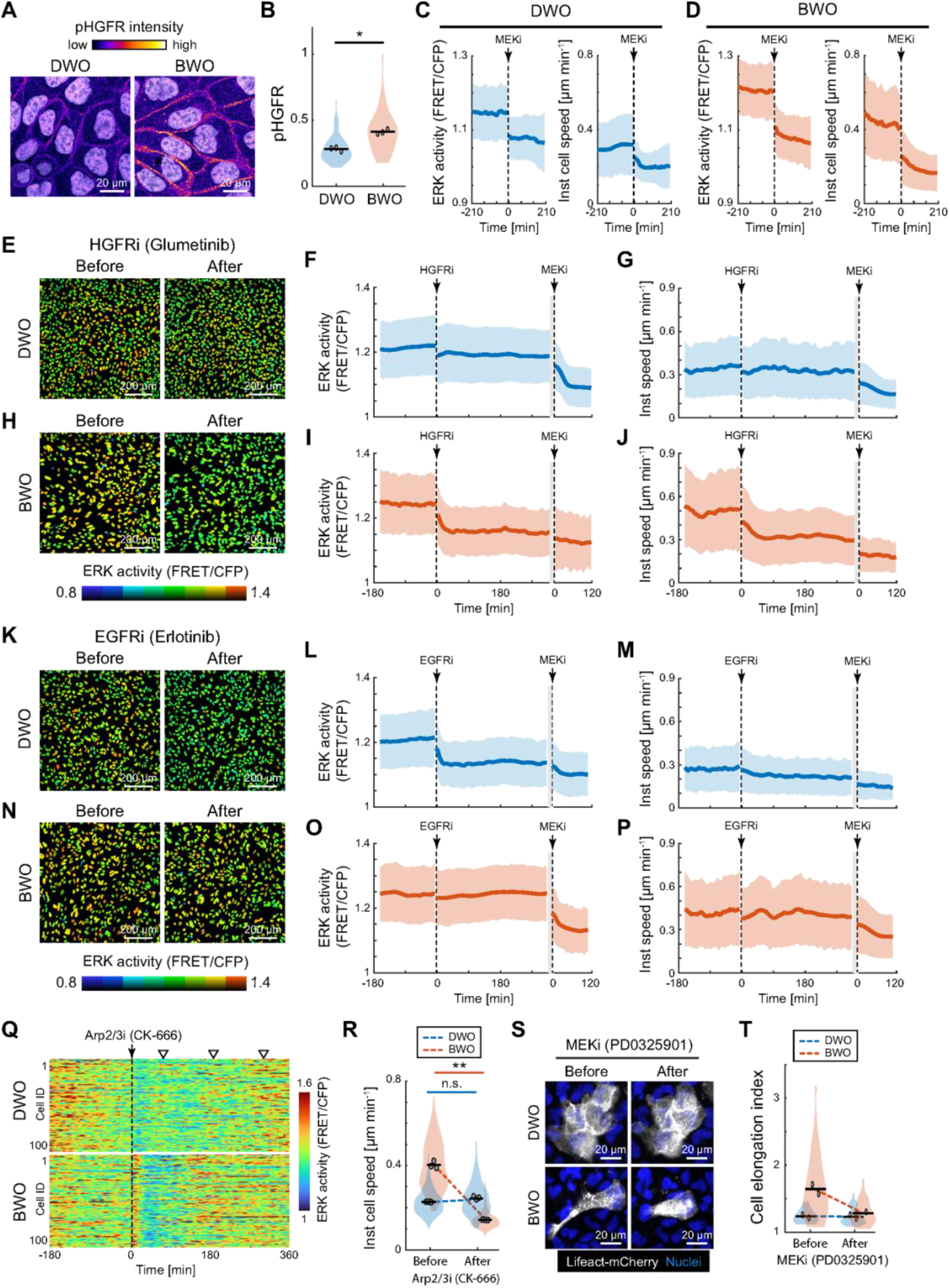
HGFR-dependent ERK signaling and actin-driven protrusion induced by BWO. (A) Immunofluorescence images of phosphorylated HGFR (pHGFR) with nuclear counterstaining (white) in DWO and BWO cells on Day 1. Scale bars, 20 µm. (B) Normalized pHGFR signal intensity at the plasma membrane, scaled to the maximum value within each dataset. DWO: n=120 cells, N=3; BWO: n=120 cells, N=3. Paired two-tailed t-test on biological-replicate means, p=0.0167. (C,D) ERK activity and instantaneous cell speed as a function of time for DWO (C) and BWO (D) cells before and after treatment with the MEK inhibitor PD0325901 (1 µM). Time zero indicates the onset of inhibitor treatment. Thick lines denote mean values, and shaded regions represent standard deviation (s.d.) across all cells. DWO: n>990 cells, N=3; BWO: n=470 cells; N=3. (E) Representative ERK activity maps of DWO monolayers before and after treatment with the HGFR inhibitor glumetinib (10 µM). Scale bar, 200 µm. (F,G) ERK activity (F) and instantaneous cell speed (G) in DWO cells as a function of time following sequential treatment with an HGFR inhibitor (glumetinib, 10 µM) and a MEK inhibitor (PD0325901, 1 µM). Time zero corresponds to the timing of each inhibitor treatment; see axis labels in panels (I) and (J). Thick vertical grey lines indicate an axis break; after the break, time was re-zeroed to align MEKi addition at t=0. Thick lines denote mean values and shaded regions indicate s.d. n>800 cells, N=3. (H) Representative ERK activity maps of BWO monolayers before and after treatment with the HGFR inhibitor glumetinib (10 µM). Scale bar, 200 µm. (I,J) ERK activity (I) and instantaneous cell speed (J) in BWO cells following sequential HGFR and MEK inhibition. Graph presentation and conventions are the same as in panel (F), unless otherwise noted. n>490 cells, N=3. (K) Representative ERK activity maps of DWO monolayers before and after treatment with the EGFR inhibitor erlotinib (500 nM). Scale bar, 200 µm. (L,M) ERK activity (L) and instantaneous cell speed (M) in DWO cells following sequential EGFR inhibition (erlotinib, 500 nM) and MEK inhibition. Graph presentation is the same as in panel (F). N>550 cells, N=3. (N) Representative ERK activity maps of BWO monolayers before and after treatment with the EGFR inhibitor erlotinib (500 nM). Scale bar, 200 µm. (O,P) ERK activity (O) and instantaneous cell speed (P) in BWO cells following sequential EGFR and MEK inhibition. Graph presentation is the same as in panel (F). n>490 cells, N=3. (Q) Heatmap of ERK activity dynamics at the single-cell level before and after treatment with the Arp2/3 inhibitor CK-666 (100 µM). Time zero indicates the onset of inhibitor treatment. One hundred representative cells were randomly selected from a local region within the time-lapse dataset. White triangles indicate periods of ERK activation in DWO cells. (R) Mean cell speed measured over 2-h windows before and 10–12 h after Arp2/3 inhibition. DWO: n=777 cells before and n=572 cells after, N=3; paired two-tailed t-test on biological-replicate mean, p=0.0543. BWO: n=572 cells before and n=639 cells after, N=3; paired two-tailed t-test on biological-replicate means, p=1.62×10^-3^. (S) Representative fluorescence images of cells in homotypic DWO–DWO co-culture (top) and heterotypic BWO–DWO co-culture (bottom) before and after treatment with the MEK inhibitor PD0325901 (1 µM). LifeAct–mCherry is shown in white and nuclei are shown in blue. Scale bars, 20 µm. (T) Cell elongation index measured before and 80 min after MEK inhibition with PD0325901 (1 µM). Dotted lines connect median values for each condition. DWO, BWO: n=30, N=2.

We next compared ERK activity and associated cell motility responses to targeted perturbations of the signaling network. Treatment with the MEK inhibitor PD0325901, which blocks ERK activation downstream of receptor signaling, robustly suppressed ERK activity and cell movement to a comparable extent in both DWO and BWO cells (**Figures 5C,D**), establishing a common basal reference for ERK downregulation. We then selectively inhibited HGFR activity using Glumetinib at 10 µM. In DWO cells, HGFR inhibition had little effect on either ERK activity levels or instantaneous cell speed (2.2% decrease and 3.8% increase, respectively; **Figures 5E–G, S5A,B; Video 5**). In contrast, in BWO cells, HGFR inhibition significantly reduced ERK activity by 6.7% and cell speed by 27.3% (**Figures 5H–J, S5A,B; Video 5**). Conversely, inhibition of EGFR activity with Erlotinib at 500 nM reduced ERK activity by 6.2% in DWO cells (**Figures 5K–M, S5C; Video 6**), consistent with previous reports that confluent MDCK cells surrounded by others at all sides rely on EGFR-dependent ERK regulation^44,52^. Strikingly, EGFR inhibition had little effect on ERK activity or instantaneous cell speed in BWO cells (2.0% and 0.5% decreases, respectively; **Figures 5N–P, S5D; Video 6**). Together, these results indicate that BWO shifts the dominant upstream control of ERK signaling from EGFR-dependent regulation toward an HGFR-dependent mode, even under fully confluent conditions.

The elevated baseline ERK activity observed in BWO cells resembles aspects of ERK dynamics reported in oncogene-activated epithelial cells, where sustained or increased baseline MAPK/ERK activity is associated with altered cell behavior and non-cell-autonomous epithelial responses^53,54^. In our system, however, this ERK-active state is initiated by transient attenuation of actomyosin contractility rather than by an introduced oncogenic mutation, and is coupled to HGFR-dependent protrusive migration. Although longer-term transcriptional or epigenetic changes may contribute to the maintenance of this state, our findings suggest that mechanical perturbation can provide a non-genetic entry point into a sustained ERK-active, protrusive epithelial program.

To determine whether this rewired ERK signaling is functionally coupled to Arp2/3-dependent protrusive activity, we inhibited the Arp2/3 complex using CK-666 at 100 µM to suppress actin-based protrusions such as lamellipodia. Upon Arp2/3 inhibition, DWO cells continued to display organized traveling waves of ERK activity, whereas BWO cells exhibited disrupted ERK activity patterns (**Figure 5Q; Video 7**). This disruption was accompanied by a significant reduction in cell speed specifically in BWO cells (**Figure 5R**). These results indicate that BWO dynamics are preferentially dependent on Arp2/3-mediated protrusive activity. The relative resistance of DWO dynamics to acute Arp2/3 inhibition suggests that this ERK-wave-associated collective mode is less dependent on Arp2/3-mediated protrusions than the BWO state, consistent with previous work distinguishing HGF–ERK–lamellipodial feedback in migratory leader cells from EGFR-dependent oscillatory ERK waves in follower cells^52^. Previous work further showed that leader-cell HGF sensitivity is promoted by lamellipodial extension and talin1-dependent immature focal complexes, rather than by increased HGFR/Met expression^52^. Thus, BWO may establish a protrusive and adhesive cellular state that favors HGFR-dependent ERK regulation. We do not infer that transient contractility attenuation directly primes HGFR through a specific molecular adaptor, and the molecular mechanism coupling these mechanical changes to altered receptor dependence remains to be determined. In BWO cells, Arp2/3-dependent protrusions may help sustain the ERK-active state. We do not infer that Arp2/3 directly activates ERK; instead, protrusions may reinforce ERK activity indirectly through HGFR/Ras/PI3K signaling, cytoskeletal remodeling, and adhesion-associated feedback. This interpretation is consistent with previous work showing that cellular protrusions can trigger ERK activation through feedback involving Ras, PI3K, the cytoskeleton, and cell adhesion^55^. Moreover, we examined single-cell morphological changes under confluent homotypic co-culture conditions, in which one cell population was fluorescently labeled. Suppression of ERK activity using MEK inhibition caused BWO cells to lose their elongated morphology and adopt a more rounded shape, with the mean elongation index decreasing from 1.65 to 1.29, whereas DWO cell morphology remained largely unchanged (1.24 to 1.23; **Figures 5S,T**). Collectively, these results demonstrate that BWO establishes an HGFR-dependent, protrusion-coupled ERK signaling state in which ERK activity is linked to actin-based protrusive activity, thereby supporting persistent collective migration.

### Protrusion-driven migration promotes shape–velocity alignment and enables coherent, persistent collective flow

To understand how protrusion-driven migration gives rise to coherent multicellular flow, we developed a two-dimensional cellular Potts model (CPM)^44,56–58^ to describe the dynamics of a two-dimensional MDCK epithelial model tissue, corresponding to a confluent monolayer, at single-cell resolution (**Figures 6A,A’**). The model incorporates minimal mechanical ingredients governing multicellular behavior, including intercellular adhesion, cell elasticity, and active migration. Cells were endowed with an intrinsic polarity vector representing front–rear asymmetry associated with either contractile or protrusive migration (**Figure 6B**). The polarity direction evolves dynamically through alignment with the resultant cell velocity arising from neighbor interactions, balanced by stochastic fluctuations. Because organized ERK–contractility feedback has been shown previously to generate pulsatile waves in epithelial monolayers^43,44,59^, our aim here was not to reconstruct the full mechanochemical dynamics of DWO cells. Instead, we used a minimal CPM to test a complementary question: whether a switch in the dominant mechanical mode of migration is sufficient to promote shape–velocity alignment and coherent persistent motion. We therefore treated contraction-driven and protrusion-driven migration as limiting regimes, rather than exact representations of DWO and BWO cellular states. Explicit incorporation of ERK–contractility feedback would be required to quantitatively reproduce the DWO wave dynamics and represents an important extension of the present minimal framework. To isolate the effects of migration mode, we considered two regimes: contraction-driven migration (*m_c_* > 0, *m_p_* = 0) and protrusion-driven migration (*m_c_* = 0, *m_p_* > 0).

**Figure 6.**
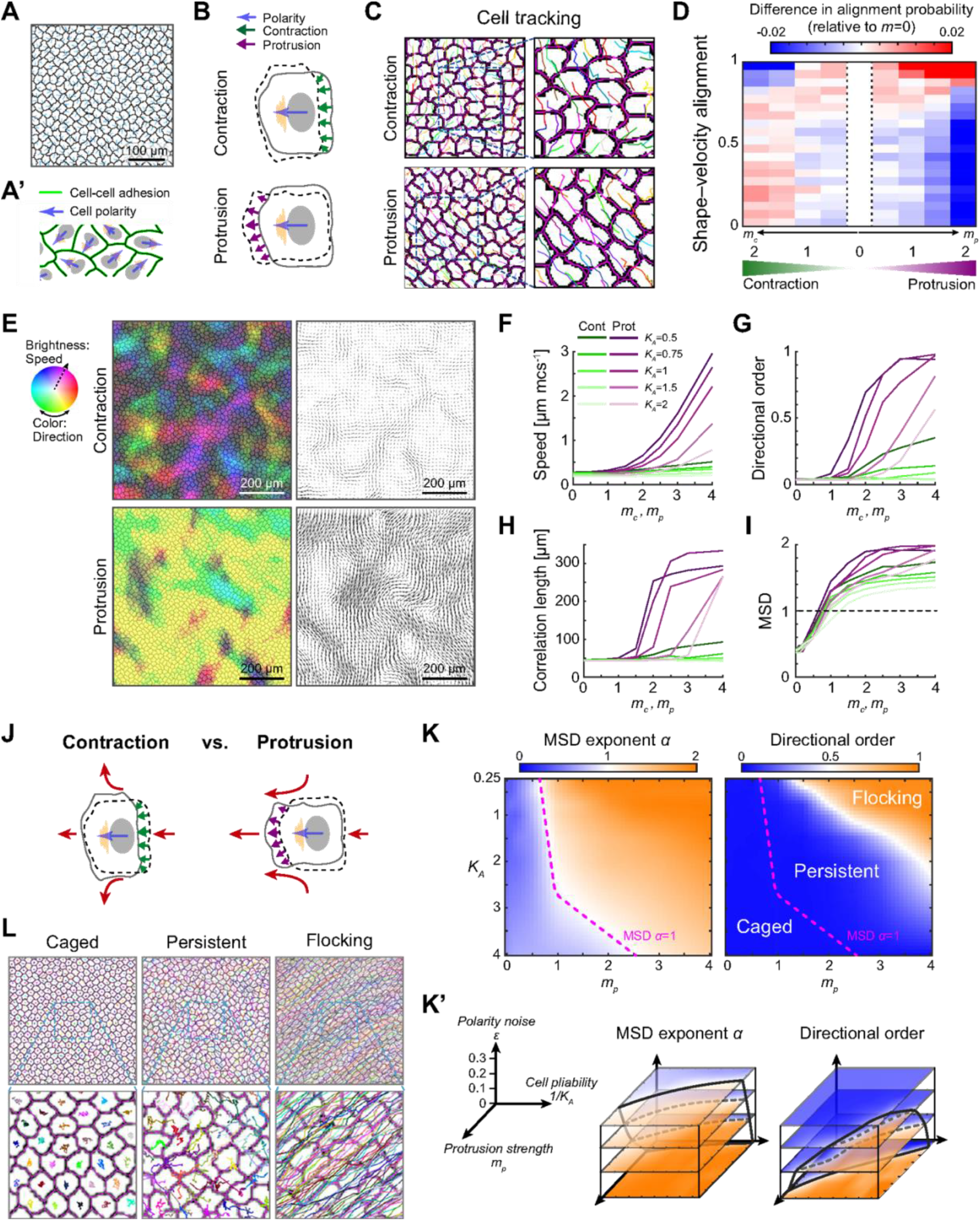
Mathematical model reveals that protrusion-driven migration promotes coherent collective flow. (A,A’) Simulated epithelial monolayer in the Cellular Potts Model (CPM) (A), in which cells adhere to one another and exhibit front-rear polarity (blue arrows). Panel (A’) illustrates cell-cell adhesion and polarity vectors. Scale bar, 100 µm. (B) Two modes of cell migration implemented in the model: rear contraction (top) and front protrusion (bottom). (C) Representative snapshots of single-cell trajectories under contraction-driven (top) and protrusion-driven (bottom) migration. Colored lines indicate individual cell trajectories. (D) Difference in shape–velocity alignment probability relative to the non-motile condition (*m_c_* = *m_p_* = 0). Parameters: *K_A_* = 1, *ε* = 0.1. (E) Representative maps of optical flow (left) and corresponding velocity vector fields (right) for contraction-driven (top) and protrusion-driven (bottom) regimes. Parameters: *m_c_* = 2 for contraction and *m_p_* = 2 for protrusion; *K_A_* = 1, *ε* = 0.2. Scale bars, 200 µm. (F–I) Dependence of instantaneous speed (F), directional order parameter (G), spatial correlation length (H), and MSD exponent *α* (I) on model parameters (*m_c_*, *m_p_*, and *K_A_*). *ε* = 0.2. Green curves indicate the contraction regime, and purple curves indicate the protrusion regime at different values of *K_A_*. (J) Schematic illustrating how distinct migration modes give rise to different collective behaviors. Red arrows indicate how cell deformation of each mode influences the motion of neighboring cells. (K,K’) Phase diagrams of MSD exponent (*α*, left) and directional order (right) as functions of protrusion strength *m_p_* and cell elasticity *K_A_* in the protrusion-driven regime, with contraction strength set to zero. The MSD map distinguishes subdiffusive (*α* < 1) and superdiffusive (*α* > 1) regimes. Combined with directional order, three behavioral regimes emerge: caged, persistent, and flocking. These phase boundaries are robust across polarity noise levels (*ε* = 0, 0.1, 0.2, 0.3), as shown in panel (K’). Black solid and dashed lines in panel (K’) correspond to boundaries identified in the 2D phase maps in panel (K). (L) Representative cell trajectories in each regime: caged (left, *m_p_* = 0.25), persistent (middle, *m_p_* = 1.5) and flocking (right, *m_p_* = 3), with *K_A_* = 1 and *ε* = 0.2.

We first examined how migration mode influences the relationship between cell shape and migration direction. In both regimes, cells exhibited elongation due to polarized activity; however, the orientation of elongation differed markedly. In the contraction-driven regime, cells elongated predominantly transverse to the migration direction (**Figure 6C, top**), consistent with behaviors observed in DWO and untreated cells (**Figure 4B, top**). In contrast, protrusion-driven migration produced elongation aligned with the migration direction (**Figure 6C, bottom**), recapitulating the behavior of BWO cells (**Figure 4B, bottom**). To quantify this relationship, we introduced a shape–velocity alignment metric, defined as the inner product between the cell elongation axis and the velocity vector. Increasing contraction strength *m_c_* decreased alignment values, indicating motion transverse to elongation. Conversely, increasing protrusion strength *m_p_* enhanced positive alignment, indicating motion along the elongated axis (**Figure 6D**). These opposing trends arise from distinct deformation mechanisms: rear contraction compresses the cell along the polarity axis, inducing lateral expansion, whereas front protrusion extends the cell along the polarity axis (**Figure 6B**).

We next investigated how these migration modes influence collective dynamics. Despite having comparable strengths (*m_c_* or *m_p_*), the contraction-driven regime produced pulsatile, spatially confined motion with limited displacement (**Figure 6E, top; Video 8**), whereas the protrusion-driven regime generated long-range, persistent flows (**Figure 6E, bottom; Video 9**). Quantitative analysis using instantaneous speed, directional order, spatial correlation length, and MSD revealed that increasing migration strength enhanced all metrics in both regimes; however, the protrusion-driven regime exhibited a markedly stronger sensitivity (**Figures 6F–I**). In addition, reducing cell elasticity *K_A_*, corresponding to increased cell deformability, substantially enhanced speed, directional order, and correlation length, with a more modest effect on MSD. These results identify migration strength and cell deformability as key control parameters governing the emergence of coherent collective motion, consistent with the experimentally observed properties of BWO cells. Mechanistically, the two migration modes generate distinct supracellular flow fields. Contraction-driven migration induces lateral displacement of neighboring cells, promoting divergence and spatially confined motion with limited net displacement. In contrast, protrusion-driven migration pulls neighboring cells inward and forward, promoting convergence and alignment at supracellular scales. This difference in mechanical coupling leads to the emergence of coherent and persistent collective flow in the protrusion-driven regime (**Figure 6J**). Thus, we do not propose that ERK directly specifies the boundaries of coherent motion domains. Rather, ERK/HGFR signaling supports the protrusive motile state, while long-range coupling can emerge physically from shape–velocity alignment and local mechanical interactions among elongated motile cells, as described in active-matter models of collective motion^60–62^.

Finally, to characterize the protrusion-driven regime relevant to BWO cells, we systematically explored the parameter space defined by protrusion strength *m_p_* and cell elasticity *K_A_* while setting contraction strength to zero (*m_c_* = 0). This analysis revealed three distinct dynamical regimes: caged, persistent, and flocking (**Figures 6K–L, S6**). At low *m_p_*, cells exhibited low persistence and low directional order, characteristic of a solid-like, caged state (**Figure 6L, left**). At intermediate *m_p_*, cells displayed superdiffusive motion with limited directional order, corresponding to a persistent but disordered regime (**Figure 6L, middle**). At high *m_p_*, both MSD and directional order increased markedly, indicating a transition to a flocking-like state with highly coordinated motion (**Figure 6L, right**). While polarity noise modulated the phase boundaries (**Figure 6K’**), the overall structure of the phase diagram remained robust. Together, these results demonstrate that protrusion-driven migration promotes shape–velocity alignment and enables coherent, persistent collective flow, while the resulting phase framework offers a basis for interpreting potentially unexplored epithelial behaviors.

Altogether, our results demonstrate that transient attenuation of actomyosin contractility is sufficient to reprogram epithelial cells into a protrusion-driven state, coordinating changes across multiple mechano-chemical layers, including cytoskeletal organization, cell–substrate adhesion, ERK signaling, and migration mode, even under confluent condition. Thus, while reduced cortical tension and increased strain may create a permissive physical state for fluidization, our results suggest that BWO actively regulates this transition by shifting epithelial cells toward a protrusion-driven, HGFR/ERK-coupled migration mode. BWO cells consequently share several morphological, adhesive, protrusive, and signaling features with previously described leader cells; however, these similarities do not establish functional equivalence to bona fide leader cells, which would require a dedicated quantitative leader–follower assay. Our findings therefore show that several leader-like features can emerge following an internal mechanical state transition, rather than being restricted to cells exposed to chemical or geometric cues at tissue boundaries. Importantly, this reprogramming occurs during the recovery phase following contractility attenuation, a period likely accompanied by extensive remodeling of protein states and gene expression programs. In line with a previous work showing that reduced intracellular tension promotes chromatin accessibility and cellular reprogramming^63^, our results raise the possibility that mechanical relaxation may couple to epigenetic regulation to stabilize this migratory state. Because the BWO protocol involves a 16-h pharmacological perturbation followed by washout and recovery, it should not be considered a direct mimic of brief physiological reductions in contractility. Rather, our findings establish that a temporally restricted attenuation of contractility followed by recovery can induce a persistent protrusion-driven migratory state. Whether analogous state transitions arise during physiological changes in contractility, such as those accompanying tissue stretching, wound repair, or morphogenesis, remains to be determined.

## MATERIALS AND METHODS

### Cell lines and culture

MDCK cells were obtained from the RIKEN BioResource Center (RCB0995). Lenti-X 293T (HEK293) cells were purchased from Takara Bio (Cat. #632180). EKAREV-NLS–expressing MDCK cells (MDCK-ERK) were established as described previously^44^. Lifeact-mCherry–expressing MDCK cells (MDCK-Lifeact-mCherry) were a gift from Michiyuki Matsuda’s laboratory. All cells were maintained in Dulbecco’s modified Eagle’s medium (DMEM; Thermo Fisher Scientific, #11965092) supplemented with 10% heat-inactivated fetal bovine serum (FBS; Thermo Fisher Scientific, #A5670801) and 1% penicillin–streptomycin (10,000 U mL⁻¹; Thermo Fisher Scientific, #15140148), hereafter referred to as the complete medium. Cultures were maintained in a humidified incubator at 37°C with 5% CO₂.

### Generation of DWO and BWO cells

DMSO-washout (DWO) and blebbistatin-washout (BWO) cells were generated following a previously described protocol ^25^. Briefly, MDCK cells (1×10⁵ cells) were seeded into each well of a 6-well plate and cultured for 20–24 h to allow cell attachment and spreading. Cells were then treated with either dimethyl sulfoxide (DMSO; Thermo Fisher Scientific, #D12345) or (−)-blebbistatin (Selleckchem, #S7099). For blebbistatin treatment, the complete medium was replaced with 2 mL of complete medium containing blebbistatin at a final concentration of 40 µM, and cells were incubated for 16 h. Control cells received an equivalent volume of DMSO. Following drug treatment, cells were washed three times with fresh complete medium and incubated in complete medium for an additional 4 h to allow recovery. Cells were then detached using 0.25% trypsin–EDTA (Thermo Fisher Scientific, #25200072) and reseeded onto the appropriate substrates coated with 20 µg mL⁻¹ type I collagen (Sigma-Aldrich, #C4243) for each experiment at either near-confluent density (1,100 cells mm⁻²) or sparse density (55 cells mm⁻²).

### Co-culture assay

MDCK–Lifeact-mCherry cells were used as either BWO or DWO populations, while MDCK-ERK cells served as the counterpart population in co-culture assays (i.e., untreated or DWO cells). Following the 4 h recovery period after washout, the two cell populations were thoroughly mixed at a 1:1 ratio for Figure 1 and at a 1:20 (label vs. non-label) ratio for Figure 5 in a tube and seeded into each well of a 24-well plate. Cells were then maintained in the complete medium, with the medium replaced daily.

### Monolayer expansion and confluent assays

For the monolayer expansion assay, MDCK cells were confined using a Culture-Insert 2 Well in µ-Dish 35 mm (ibidi, #81176). MDCK cells (2×10⁴ cells per well) were seeded into each compartment of the insert and cultured for 24 h in the complete medium to allow the formation of confluent monolayers. Confinement was released by manual removal of the insert, and the medium was immediately replaced with FluoroBrite DMEM (Thermo Fisher Scientific, #1896701) supplemented with 10% fetal bovine serum (FBS) and 1% penicillin–streptomycin, hereafter referred to as the imaging medium. For confluent monolayer assays, MDCK cells were seeded at near-confluent density onto collagen-coated glass-bottom dishes (Iwaki, #3971-035) and cultured for 24 h in the complete medium until full confluence was reached. The medium was then replaced with the imaging medium. For both conditions, time-lapse imaging was performed using a Nikon AX R confocal microscope equipped with a CFI Plan Fluor DLL 20× objective (NA = 0.5), with images acquired at 5-min intervals. For FRET imaging, donor excitation was performed using a 445 nm laser. Emission was collected using band-pass filters of 464–499 nm for the CFP (donor) channel and 518–551 nm for the FRET (acceptor) channel. Images were acquired using galvano scanning with 2× line averaging. For FRET image analysis, raw images were denoised using a 3 × 3 median filter, and background fluorescence was subtracted independently from the CFP and FRET channels. FRET/CFP ratio images were then generated using a custom MATLAB (MathWorks) script. In the resulting pseudocolor images, hue represents the FRET/CFP ratio, while brightness corresponds to the intensity of the FRET channel.

### Antibodies, dyes, and inhibitors

The following chemicals were used: Mouse monoclonal anti-phospho-p53 (Cell Signaling Technology, #9286; 1:200 dilution), Mouse monoclonal anti-p21 (Santa Cruz, #sc-6246, 1:200 dilution), Rabbit monoclonal anti–E-cadherin (Cell Signaling Technology, #3195; 1:200 dilution), Rabbit monoclonal anti-Paxillin (Abcam, #ab32084; 1:500 dilution), Rabbit monoclonal anti-phospho-Met (Tyr1234/1235) (Cell Signaling Technology, #3077; 1:100), Goat anti-mouse IgG (H+L) Alexa Fluor 647 (Abcam, #ab150115; 1:1000 dilution), Goat anti-rabbit IgG (H+L) Alexa Fluor 546 (ThermoFisher, #A-11035; 1:1000 dilution), Goat anti-rabbit IgG (H+L) Alexa Fluor 647 (Abcam, #ab150079; 1:1000 dilution), Hoechst 33342 (Thermo Fisher Scientific, #62249; 1:1000 dilution), Alexa Fluor™ 647 phalloidin (Invitrogen, #A22287; 1:500 dilution), Glumetinib (MedChemExpress, #HY-116000), Erlotinib (MedChemExpress, #HY-50896), CK-666 (Merck, Sigma-Aldrich, #SML0006), PD0325901 (Sigma-Aldrich, #PZ0162).

### Immunofluorescence staining and imaging

For immunofluorescence staining, cells were fixed with 4% paraformaldehyde (PFA) for 15 min at room temperature (23°C) and washed twice with phosphate-buffered saline (PBS) for 5 min each. Fixed cells were permeabilized with 0.1% Triton X-100 in PBS (PBST) for 5 min and subsequently washed with PBS. Samples were then blocked with 2% normal goat serum (Cell Signaling Technology, #5425S) for 2 h at 37°C. Cells were incubated with primary antibodies diluted in PBS overnight at 4°C. After three washes with PBS, samples were incubated with appropriate secondary antibodies and/or fluorescent staining dyes for 2 h at 37°C or overnight at 4°C. Following staining, samples were washed twice with PBS and stored in PBS at 4°C until imaging. Fluorescence images were acquired using an AX R confocal microscope (Nikon) equipped with a CFI Plan Apochromat VC 60× water-immersion objective (NA = 1.2) or a CFI Plan Apochromat Lambda S 40× silicone-immersion objective (NA = 1.25).

### Centrifugation assay and analysis

For the centrifugation assay, HEK293 cells were used instead of MDCK cells, as MDCK monolayers did not detach even at the maximum centrifugal force attainable with our equipment. DWO- or BWO-treated HEK293 cells were reseeded at a density of 100 cells mm⁻² onto collagen-coated circular glass coverslips (18 mm diameter) and cultured overnight in complete medium to allow cell adhesion. Each coverslip was placed into a custom-designed centrifugation stage, which was inserted into a 50-ml Falcon tube containing 10 ml of complete medium. Samples were centrifuged using a Sorvall X Pro centrifuge (Thermo Fisher Scientific) for 3 min at 37°C. Following centrifugation, cells remaining on the coverslip were fixed with 4% PFA for 15 min, washed three times with PBS each 5 min each, and mounted onto glass microscope slides using Fluoromount™ Aqueous Mounting Medium (Sigma-Aldrich, #F4680). Imaging was performed using a Nikon AX R confocal microscope equipped with a CFI Plan Apochromat Lambda 10× objective (NA = 0.45). Mechanical loading during centrifugation was quantified as the effective centrifugal force acting on individual adherent cells. The applied centrifugal force per cell *F* was estimated as: *F* = (*ρ_c_* − *ρ_m_*)*V* ⋅ *g* ⋅ *RCF*, where *ρ_c_* is the density of the cell (1.078 g cm⁻³), *ρ_m_* is the density of the culture medium (assumed to be 1.0 g cm⁻³), *V* is the cell volume (3 pL), *g* is the gravitational acceleration (9.81 m s⁻²), and *RCF* is the relative centrifugal force applied.

### FRAP assay and analysis

Fluorescence recovery after photobleaching (FRAP) experiments were performed using wild-type MDCK cells transiently transfected with Paxillin–pEGFP (Addgene, plasmid #15233) using Lipofectamine LTX (Thermo Fisher Scientific, #15338100) according to the manufacturer’s instructions. Twenty-four hours after transfection, cells were subjected to either DWO or BWO treatment as described above and then seeded under confluent conditions onto glass-bottom dishes (Iwaki, #3971-035). FRAP imaging was carried out using a Nikon Eclipse Ti-2 AX R confocal microscope. Imaging was performed with a CFI Plan Apochromat VC 60× water-immersion objective (NA = 1.2), using 20× digital zoom, the maximum pinhole size, and a 500 ms frame interval, at a spatial resolution of 128 × 128 pixels. A circular region of interest (ROI) with a diameter of 1 µm was selected for photobleaching. Time-lapse images were acquired for 1 min before bleaching using a 488 nm laser at 1% power to establish baseline fluorescence. Photobleaching was performed for 5 s using the 488 nm laser at 100% power, followed by post-bleach imaging with the 488 nm laser at 1% power. Fluorescence recovery curves were fitted with a single-exponential function, and the half-recovery time was determined from the fitted curve.

### Traction force microscopy

Traction force microscopy (TFM) was performed as previously described with slight modifications^44^. Briefly, MDCK cells were cultured in imaging medium on collagen-coated polyacrylamide (PA) gels with an elastic modulus of 9 kPa, embedded with fluorescent beads (0.2 µm diameter; Thermo Fisher Scientific, #F8807). Cells were plated at the near-confluent density, and live-cell imaging was performed to quantify the beads displacement generated by collective cell behavior. Polyacrylamide gel solutions containing fluorescent beads were prepared by vortex mixing 373.5 µL 1× PBS, 93.75 µL of 40% acrylamide, 25 µL of 2% bis-acrylamide, 3.2 µL fluorescent beads, 0.25 µL TEMED, and 2.5 µL of 10% ammonium persulfate. To prepare silanized coverslips, glass coverslips (22 × 22 mm) were cleaned by immersion in 1% Hellmanex III Cleaner (Sigma-Aldrich, #Z805939) for 10 min, rinsed three times with distilled water, soaked in warm distilled water at 42 °C for 60 min, and dried using pressurized nitrogen gas. Coverslips were then silanized in a solution containing 2% (v/v) 3-(trimethoxysilyl)propyl methacrylate (Sigma-Aldrich, #440159) and 1% (v/v) glacial acetic acid (Sigma-Aldrich, #A6283) in absolute ethanol for 10 min. After silanization, coverslips were washed three times with absolute ethanol (2 min each), baked at 80°C for 2 h, cooled to room temperature, and stored in an airtight container until use. Then, a 50 µL aliquot of the gel solution was pipetted onto a hydrophobic glass slide coated with Rain-X, and the silanized coverslip was gently placed on top. Gels were allowed to polymerize on a level surface for 30 min, after which the coverslip bearing the PA gel was carefully detached and soaked in distilled water overnight. For surface functionalization, Sulfo-SANPAH (Sigma-Aldrich, #80332) was dissolved in anhydrous DMSO to a concentration of 0.02 mg mL⁻¹. Two microliters of this stock solution were diluted in 40 µL of cold 20 mM acetic acid, and 42 µL of the diluted solution was applied to each PA gel. Gels were exposed to UV light for 5 min at 6% power (10 mW cm⁻²) to activate crosslinking. This activation step was repeated once using a fresh Sulfo-SANPAH solution, followed by a wash with cold 20 mM acetic acid for 5 min. Type I collagen was diluted in cold 20 mM acetic acid to a final concentration of 20 µg mL⁻¹, and gels were coated with 2 mL of the collagen solution overnight at 4 °C, after which they were washed with 1× PBS at room temperature and stored submerged in PBS until cell seeding. Gel-bearing coverslips were transferred to stainless steel cell culture chambers (Aireka Scientific, #SC15022), and 500 µL of the imaging medium was added to maintain cell confluence. Live-cell imaging was performed using a Nikon AX R confocal microscope equipped with a CFI Plan Fluor DLL 20× objective (NA = 0.5), with images acquired at 5-min intervals. Following time-lapse imaging, cells were removed by trypsinization, and reference images of the relaxed bead positions were obtained. Traction force reconstruction was performed using the u-interforce MATLAB package developed by the Danuser laboratory^64^. Bead displacement fields were calculated by comparing stressed and reference images, and traction stresses were computed according to the standard workflow implemented in the software.

### Cell tracking

Nuclear segmentation and single-cell tracking were performed using the TrackMate–Cellpose plugin implemented in ImageJ/Fiji (National Institutes of Health). For preprocessing, the FRET-channel images acquired from live imaging of MDCK-ERK cells were used as input. Images were first denoised using a 3 × 3 median filter to reduce shot noise. Pixel intensities were then normalized using contrast-limited adaptive histogram equalization (CLAHE) with default ImageJ parameters to enhance nuclear contrast. Nuclei were detected using the Cellpose detector within TrackMate. The target object diameter was set in the range of 15–25 µm, depending on the sample and imaging conditions. Detected nuclei were subsequently linked over time using TrackMate’s built-in tracking algorithms with default linking and gap-closing parameters. The resulting trajectories were used for downstream quantitative analyses.

### Optical flow analysis

Optical flow analysis was performed in MATLAB (MathWorks) using the Farnebäck dense-flow algorithm to quantify pixel-wise motion of cell collectives. Time-lapse image sequences were first registered to correct for sample drift. Dense optical flow fields were then computed for each pair of consecutive frames using the opticalFlowFarneback function from the Computer Vision Toolbox, with parameter settings optimized for the spatial scale and velocity range characteristic of the system. The resulting optical flow vectors were visualized either as instantaneous velocity arrows or as color-coded speed maps or were used for spatial correlation analysis. In velocity-direction maps, color indicates the direction of motion and brightness indicates speed. The center of the color wheel is transparent and denotes zero or undefined velocity direction. All analyses and visualizations were performed using custom MATLAB scripts.

### Density-matched analysis

Nuclear density was calculated as the number of detected nuclei through the TrackMate–Cellpose divided by the imaged area and expressed as cells/mm². To compare DWO and BWO monolayers over overlapping density ranges, DWO cells were additionally seeded at lower densities. Because BWO monolayers exhibited spatial heterogeneity in cell density within the original 1.38 -mm field of view (FOV), smaller local regions (513-µm FOV) with relatively homogeneous nuclear density throughout the 150-min observation period were selected for analysis. Instantaneous cell speed, directional order, and MSD were quantified within these local regions using the same procedures as for the corresponding whole-field analyses.

### Cross-correlation analysis

Single-cell cross-correlation analysis was performed in MATLAB using custom scripts and built-in functions. ERK activity and instantaneous cell speed were obtained from the same single -cell tracking data. Only cells continuously tracked over the analyzed time window were included. ERK activity and speed time series were aligned for each cell, and Pearson correlation coefficients were calculated at temporal lags from −120 to +120 min using MATLAB’s built-in corr function. Population-averaged cross-correlation curves were calculated by averaging single-cell cross-correlation coefficients at each lag using mean. For quality-control filtering, cells with low ERK or speed variability and cells containing extreme ERK or speed outliers were excluded using predefined thresholds based on the MATLAB functions, such as std, quantile, prctile, and median absolute deviation (mad).

### Quantification of cell area and elongation

Phase-contrast images of confluent MDCK monolayers and isolated single cells were used for morphological analysis. Cell boundaries were identified by thresholding in Fiji/ImageJ, and each enclosed region was analyzed as an individual particle using the Analyze Particles function. Cell area and circularity were measured for each cell. The cell elongation index was calculated as the inverse of circularity, with larger values indicating greater elongation.

### Quantification of junctional E-cadherin and cortical actin intensity

E-cadherin and cortical actin intensities were quantified using Fiji/ImageJ. For each image, linear regions of interest (ROIs) were manually traced along continuous apical junctional/cortical signals. For E-cadherin staining, ROIs were drawn along E-cadherin-positive cell–cell junctions. For phalloidin staining, ROIs were drawn along the apical cortical F-actin signal at cell–cell junctions or cell edges. Mean fluorescence intensity within each ROI was measured using the Fiji “Measure” function. The analysis was therefore restricted to junctional/cortical signal intensity and did not quantify total cellular phalloidin intensity. ROI length and mean intensity values were exported from Fiji for subsequent analysis.

### Quantification of phospho-p53 and p21 nuclear enrichment

Immunofluorescence staining for phospho-p53 and p21 was performed as described above. DWO and BWO cells expressing EKAREV-NLS were imaged under confluent conditions, and the EKAREV-NLS signal was used to define nuclear regions. Nuclei were segmented using a custom MATLAB pipeline incorporating background correction, adaptive thresholding, morphological filtering, and watershed separation of touching nuclei, followed by size and shape filtering. For each segmented nucleus, a perinuclear cytoplasmic annulus was defined between 3 and 10 pixels outside the nuclear boundary, with pixels belonging to neighboring nuclei excluded. For each nucleus, the mean phospho-p53 or p21 fluorescence intensity within the nuclear region (S_n) was measured together with the mean intensity within a surrounding perinuclear cytoplasmic annulus (S_c). Nuclear enrichment was quantified as the nuclear-to-cytoplasmic intensity ratio, S_n/c = S_n/S_c.

### Quantification of focal adhesions

Individual cell boundaries were identified from cortical actin staining, and cell ROIs were manually defined. Paxillin-positive focal adhesions were segmented using a fixed intensity threshold established from DWO control cells and applied consistently to all images. Individual focal adhesion area was measured from the segmented paxillin-positive regions. Focal adhesion occupancy was calculated as the total paxillin-positive area divided by the total area of the corresponding cell ROI.

### Quantification of Trypsin-detachment assay and analysis

MDCK-ERK cells under DWO or BWO conditions were exposed to 0.25% trypsin–EDTA and imaged by time-lapse microscopy. The EKAREV-NLS nuclear signal was used to monitor cell detachment. For automated detection, the signal trace was smoothed over three frames, the pre-detachment level was defined as the median signal over the first 25% of the movie, and the post-detachment level as the median of the lowest 20% of subsequent values. Detachment time was defined as the first time point at which the signal crossed the midpoint between these two levels. When a clear automated drop was not detected, the detachment time was determined manually from the time-lapse images.

### Quantification of cell deformation during monolayer expansion

Individual cells located in the central region of expanding monolayers were manually tracked from the onset of monolayer expansion using time-lapse images acquired at 5-min intervals. For each cell, the frame at which maximal stretch occurred was identified manually from the time-lapse sequence. Because deformation propagated non-uniformly from the leading edge toward the monolayer interior, the maximal-stretch time point was determined independently for each cell. Cell boundaries were manually outlined in Fiji (ImageJ), and cell area and aspect ratio were obtained using the Measure function at the onset of expansion and at the maximal-stretch time point.

### Quantification of directional order

Directional order parameter *P* was defined as 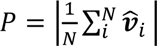, where 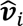 is the unit velocity vector of cell *i* and *N* is the number of cells.

### Quantification of spatial correlation

Spatial correlation function *φ* was calculated as

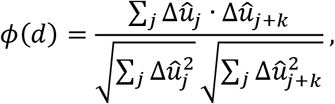

where 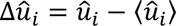 denotes the deviation of the local velocity vector 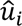 from its spatial mean, and the index *k* represents a spatial offset. The spatial index *k* was converted to the corresponding radial distance *d*. The spatial correlation length was defined as the distance at which the correlation function decayed to half of its maximum value.

### Quantification of mean squared displacement (MSD)

MSD was calculated to quantify the persistence and diffusive properties of single-cell trajectories. Cell positions were obtained from time-lapse imaging and tracked over time to generate trajectories ***r***(*t*). The MSD was defined as:

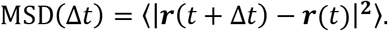

The angle brackets denote averaging over all cells and time points. MSD curves were computed over a range of time lags and plotted on a log–log scale. To characterize the mode of cell motion, MSD curves were fitted to a power-law relation: MSD(Δ*t*)∼(Δ*t*)*^α^*, where *α* is the scaling exponent. *α* = 1 indicates diffusive motion, *α* > 1 indicates superdiffusive (persistent) motion, and *α* < 1 indicates subdiffusive behavior. The exponent *α* was estimated by linear regression of the log–log MSD curves over specified time intervals.

### Quantification of cell-neighbor rearrangements

Cell-neighbor topology was quantified from cell-centroid trajectories using Delaunay triangulation at each time point in custom MATLAB scripts. Only trajectories present throughout the full imaging period were included to avoid apparent neighbor changes caused by track appearance or loss. Cells on the convex hull and their immediate Delaunay neighbors were excluded to minimize boundary effects. Neighbor retention was calculated as the fraction of a cell’s Delaunay neighbors at time *t* that remained neighbors after a given time lag and was averaged across cells and starting times. Neighbor loss represented the fraction of initial neighbors that were lost over a given time lag. To quantify local topological rearrangements, persistent Delaunay edge flips were used as a centroid-based proxy for T1-like neighbor exchanges. An event was identified when a Delaunay edge between one cell pair disappeared while the opposite diagonal within the corresponding four-cell configuration appeared. To suppress transient triangulation changes caused by positional fluctuations, the original topology was required to persist for three frames before the transition and the new topology for three frames after the transition (15 min on each side). Event rates were normalized to the number of interior cells and observation time and expressed as events per 100 cells per hour.

### Model assumptions

The cellular Potts model (CPM) was designed as a minimal model to test how different mechanical modes of migration affect collective epithelial dynamics. The model makes several simplifying assumptions. First, the epithelial monolayer is represented as a two-dimensional confluent cell sheet with periodic boundary conditions. Second, individual cells are described by area, perimeter, adhesion, and polarity-dependent migration terms, without explicitly modeling subcellular actomyosin networks, focal adhesions, or biochemical signaling pathways such as ERK. Third, contraction-driven and protrusion-driven migration are treated as limiting regimes, rather than complete representations of DWO and BWO cells. Fourth, cell polarity is updated from cell-centroid displacement with stochastic noise, allowing directional coordination to emerge from mechanical interactions rather than from an imposed neighbor-alignment rule. Thus, the model is intended to test whether protrusion-driven shape–velocity alignment is sufficient to generate coherent persistent motion, rather than to reproduce all molecular and topological features of BWO monolayers.

### Cellular Potts model

We modeled collective cell dynamics using a two-dimensional CPM, a lattice-based framework widely used to simulate multicellular systems^56–58^. In this model, each cell is represented as a connected set of lattice sites sharing an identical index *σ*. Cell behaviors emerge from stochastic updates that minimize a global effective energy ℋ. The system energy consisted of four contributions representing interfacial energy, cell area constraint, perimeter constraint, and a migration term:

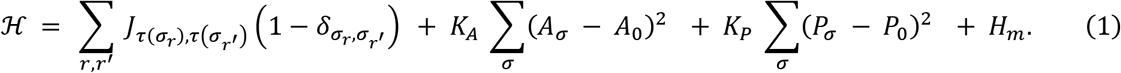

Here *r* and *r*^′^ denote neighboring lattice sites and *τ* specifies the lattice type (e.g., cell or medium). *J* represents interfacial energy between neighboring lattice identities and, *δ* is the Kronecker delta. *A_σ_* and *P_σ_* denote the current area and perimeter of cell *σ*, while *A*_0_ and *P*_0_ represent their target values. Cell perimeter was measured by raw pixel-interface counting using a Moore 8-neighbor boundary definition on the square lattice without geometric interpolation; therefore, the shape index *ρ* should be interpreted as a lattice-defined model parameter rather than a direct continuum estimate of geometric cell shape. Parameters *K_A_* and *K_P_* determine the resistance to deformation through area and perimeter constraints, respectively.

To incorporate active cell migration, we introduced an additional migration energy term

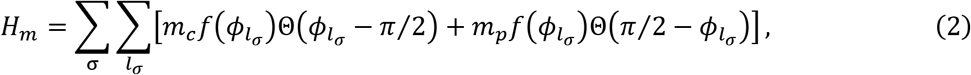

where *l_σ_* denotes lattice sites located on the cell periphery. The angle *φ_lσ_* ∈ [0, *π*] is defined between the polarity vector ***p_σ_*** and the vector connecting the cell centroid to the peripheral lattice site. The parameters *m_c_* and *m_p_* determine the strengths of contractile and protrusive activities, respectively.Θ is the Heaviside step function and *f*(*φ_lσ_*) = 2*φ_lσ_*⁄*π* − 1 modulates the angular dependence of these activities.

The dynamics of front–rear polarity was adopted from earlier studies^22,36,65,66^:

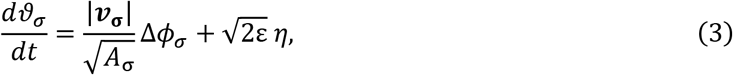

where |***v*_σ_**| is the magnitude of the velocity of cell *σ* and Δ*φ_σ_* is the angular difference between the velocity vector and the polarity vector. The parameter ε controls the strength of rotational noise, and *η* is Gaussian white noise. The first term promotes alignment between cell polarity and the direction of motion, whereas the second term introduces stochastic fluctuations.

In the CPM implementation, the polarity dynamics described above were updated in vector form. For each cell σ, the centroid displacement Δ***r***_σ_ was calculated from the change in cell centroid position between successive polarity updates, with periodic boundary correction. The polarity vector ***p***_σ_ was then biased toward the normalized displacement direction Δ***r***_σ_/|Δ***r***_σ_|, with an alignment strength proportional to 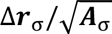, where ***A***_σ_ is the cell area. Stochastic angular noise was added to the polarity vector, and the updated polarity vector was then normalized to unit length. Thus, the factor 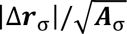 implements the velocity/displacement-dependent polarity alignment used in the simulations.

### Numerical simulations

Simulations were performed using the standard Monte Carlo dynamics of the CPM. At each update attempt, a lattice site *r* is randomly selected and its index *σ****_r_*** is proposed to be replaced by that of a randomly chosen neighboring site *σ****_r_***_′_. The change in system energy Δℋ associated with this copy attempt is evaluated. The update is accepted deterministically if Δℋ ≤ 0 and accepted with probability *exp*(−Δℋ) if Δℋ > 0. The simulation domain consisted of a square lattice of 320 × 320 pixels, corresponding to 800 × 800 µm, assuming a lattice spacing of 2.5 µm per pixel. A cluster of 1600 cells was initially placed in the domain without gaps. Periodic boundary conditions were applied in both spatial directions. All quantitative indices, including instantaneous speed, spatial correlation length, directional order, and mean squared displacement (MSD), were evaluated after transient dynamics had subsided and the system reached a quasi-steady regime in which macroscopic statistical properties remained approximately stable.

For model parameters, the target cell area *A*_0_ was set to 400 µm^2^ based on experimental measurements. The cell–cell interfacial energy *J* and the perimeter elasticity coefficient *K_P_* were fixed at 2 and 0.01, respectively, because these parameters affect the system only in a relative manner. All other parameters were varied systematically for numerical investigations. To examine the dependence on cell shape, we varied the target perimeter *P*_0_ through the shape index 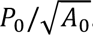. The shape index was explored within the range 3.5–4.3, which spans the solid–fluid transition regime previously reported for tissues exhibiting random cell motion in the absence of polarity^16,17^. In our model, however, polarity-dependent migration strongly deforms cell shapes, making the influence of the intrinsic shape index comparatively minor. Consequently, variations in *ρ* produced no substantial qualitative differences in collective dynamics. Unless otherwise noted, simulations were therefore performed with *ρ*=4.

### Statistical hypothesis testing

The number of cells analyzed (n) and the number of biological replicates (N) are reported in the figure legends. No statistical methods were used to predetermine sample sizes, and no randomization was performed. Statistical tests, sample sizes, test statistics, and corresponding p-values are described in the figure legends. We considered p<0.05 statistically significant for two-tailed tests and classified significance levels as follows: * (p<0.05), ** (p<0.01), *** (p<0.001), and n.s. (not significant; p≥0.05).

### Software and Graph

Digital image processing, quantitative analysis, and visualization were conducted using MATLAB R2022b, and R2024b (MathWorks) and ImageJ (National Institutes of Health). Truncated violin plots were generated using a MATLAB function sourced from MATLAB File Exchange^67^.

## ACKNOWLEDGEMENTS

This work was supported by Singapore Ministry of Education (MOE) Academic Research Fund (AcRF) Tier 2 (MOE-T2EP30223–0010), the National Research Foundation, Singapore (NRF) under its Mid-sized Grant (NRF-MSG-2023–0001), and the Mechanobiology Institute (MBI) at the National University of Singapore funded through the National Research Foundation, Singapore and the Ministry of Education, Singapore under the Research Centre of Excellence programme. We thank Pakorn Kanchanawong for providing the Paxillin-pEGFP plasmid, Chaoyu Fu and Yan Jie for insightful discussions on force measurement, and our lab members for their valuable comments on the manuscript.

## COMPETING INTERESTS

The authors declare no competing interests.

## DATA AVAILABILITY

Data collected for this study will be made publicly available via the digital institutional repository of the National University of Singapore.

## CODE AVAILABILITY

Code used for the computational simulations and analyses is available through Code Ocean.

## AUTHOR CONTRIBUTIONS

Conceptualization: TH

Methodology: SWP, TPN, TH

Investigation: SWP, SL, NN, PKM, DP, ASG, CMDM, TH

Data curation: SWP, DP, TH

Formal analysis: SWP, DP, TH

Writing – original draft: SWP, TH

Writing – review & editing: SWP, TH

Supervision: TH

Funding acquisition: TH

Resources: TH

Visualization: SWP, TH

## SUPPLEMENTARY FIGURE LEGENDS

**Figure S1.**
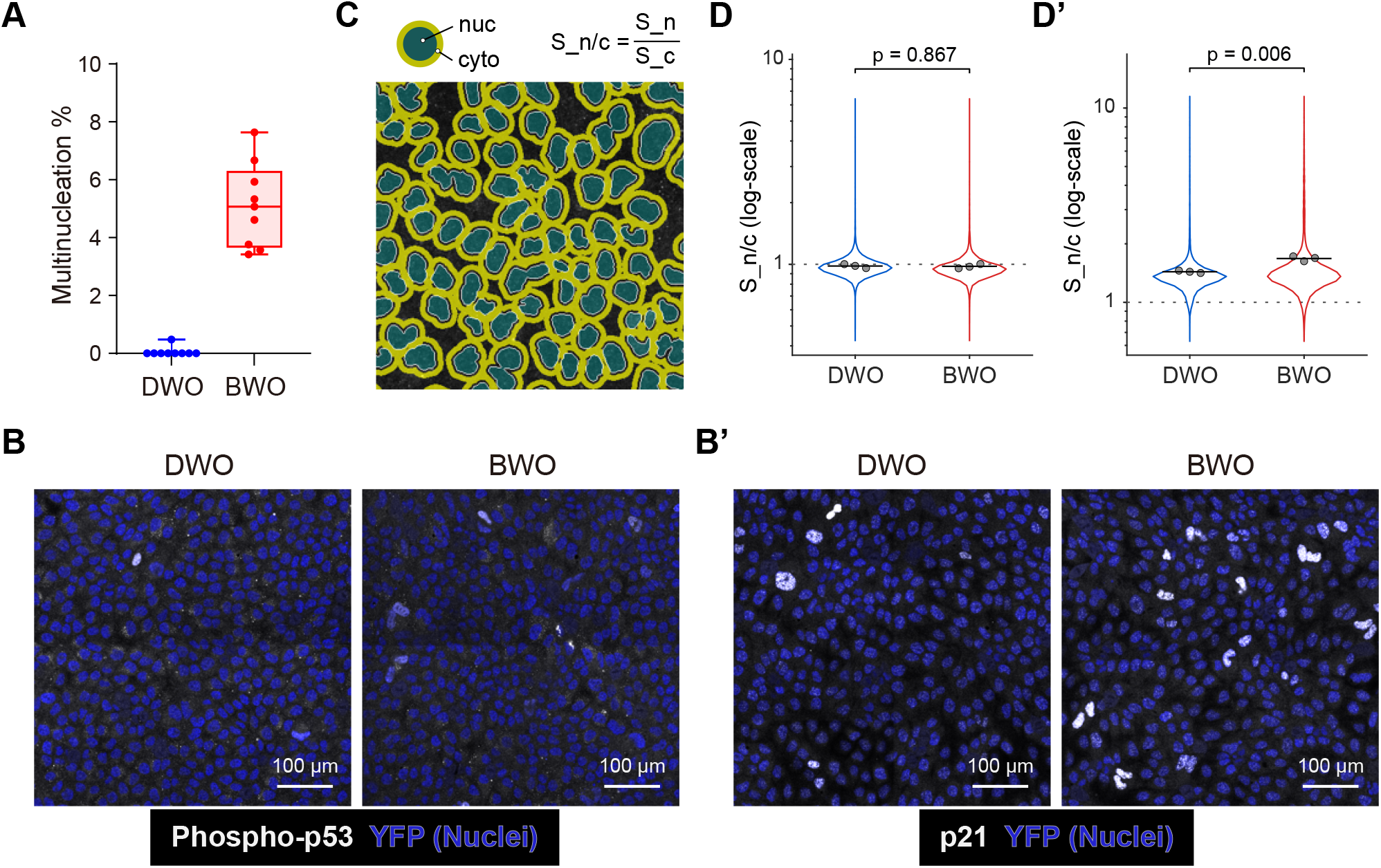
Characterization of multinucleation and p53/p21 signaling. (A) Percentage of multinucleated cells in DWO and BWO populations. Each point represents one analyzed region. DWO: 2,084 cells from nine regions, N=3; BWO: 1,266 cells from nine regions, N=3. Mean multinucleation frequency: 0.05% for DWO and 5.11% for BWO. (B,B’) Representative immunofluorescence images of phospho-p53 (B) and p21 (B’) in DWO and BWO cells. Phospho-p53 and p21 are shown in white, and the nuclear signal of EKAREV-NLS in blue. Scale bars, 100 µm. (C) Schematic of the image-analysis procedure used to quantify nuclear enrichment of phospho-p53 and p21. For each nucleus, the mean signal within the nuclear region S_n was divided by the mean signal within the surrounding perinuclear cytoplasmic annulus S_c to obtain the nuclear-to-cytoplasmic intensity ratio, S_n/c. (D,D’) Nuclear-to-cytoplasmic intensity ratios of phospho-p53 (D) and p21 (D’) in DWO and BWO cells. Violin plots show the distributions of individual nuclei; open circles indicate the means of three analyzed regions, and horizontal black lines indicate the mean of the region means. (D) Phospho-p53: DWO, n=8,564 nuclei; BWO, n=7,023 nuclei; three regions per condition. Welch’s two-sample t-test on region means, p=0.867. (D’) p21: DWO, n=7,514 nuclei; BWO, n=6,424 nuclei; three regions per condition. Welch’s two-sample t-test on region means, p=0.0062.

**Figure S2.**
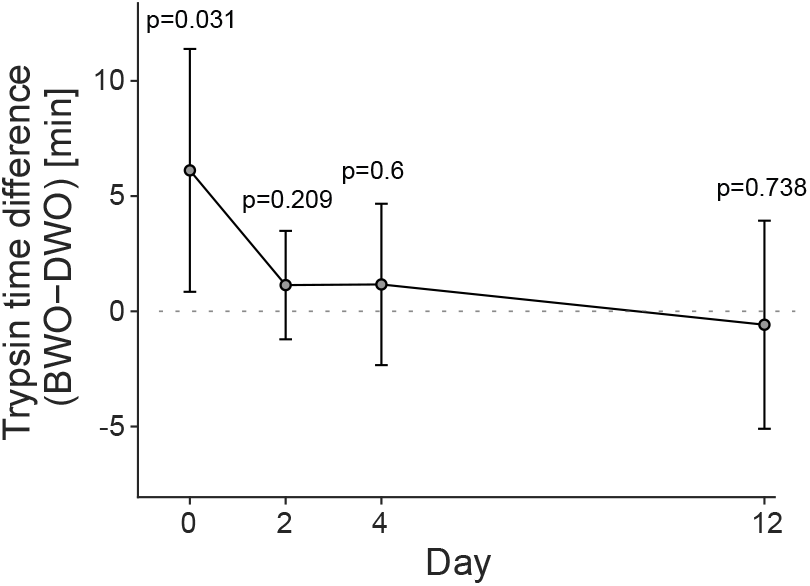
Difference in trypsin-induced cell detachment time between BWO and DWO cells over time. Difference in trypsin-induced cell detachment time between BWO and DWO cells measured at the indicated days. Values represent the difference between the mean BWO and DWO detachment times (BWO−DWO). Error bars indicate 95% confidence intervals calculated using Welch’s unequal-variance method. BWO cells showed a longer detachment time than DWO cells at Day 0 (mean difference, 6.12 min; 95% CI, 0.85–11.39 min; Welch’s t-test, p = 0.031), whereas no significant differences were detected at Day 2 (p = 0.209), Day 4 (p = 0.600), or Day 12 (p = 0.738). n≥3 per condition. The dotted line indicates no difference between BWO and DWO cells.

**Figure S3.**
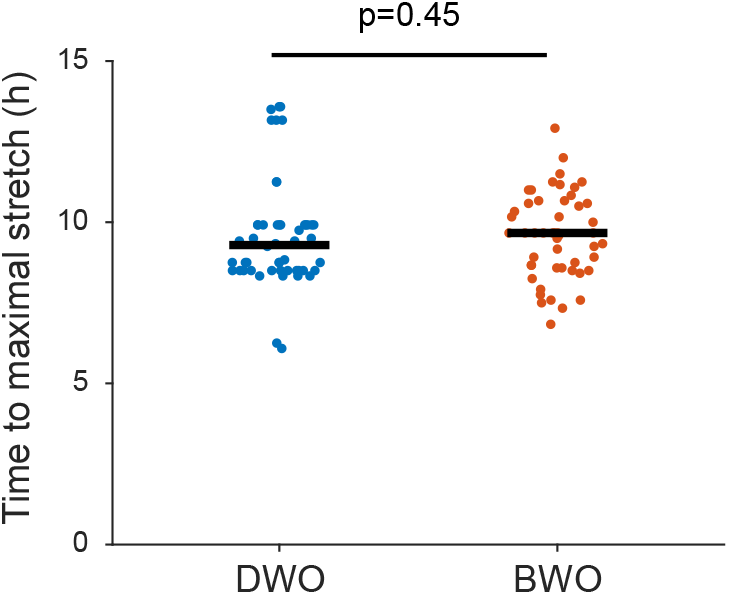
Time to maximal cell stretch during monolayer expansion. Time to maximal cell stretch in DWO and BWO cells during monolayer expansion. Each point represents one cell, and black horizontal lines indicate the median. n=50 cells, N=2; Mann–**Whitney U test, p=0.45.**

**Figure S4.**
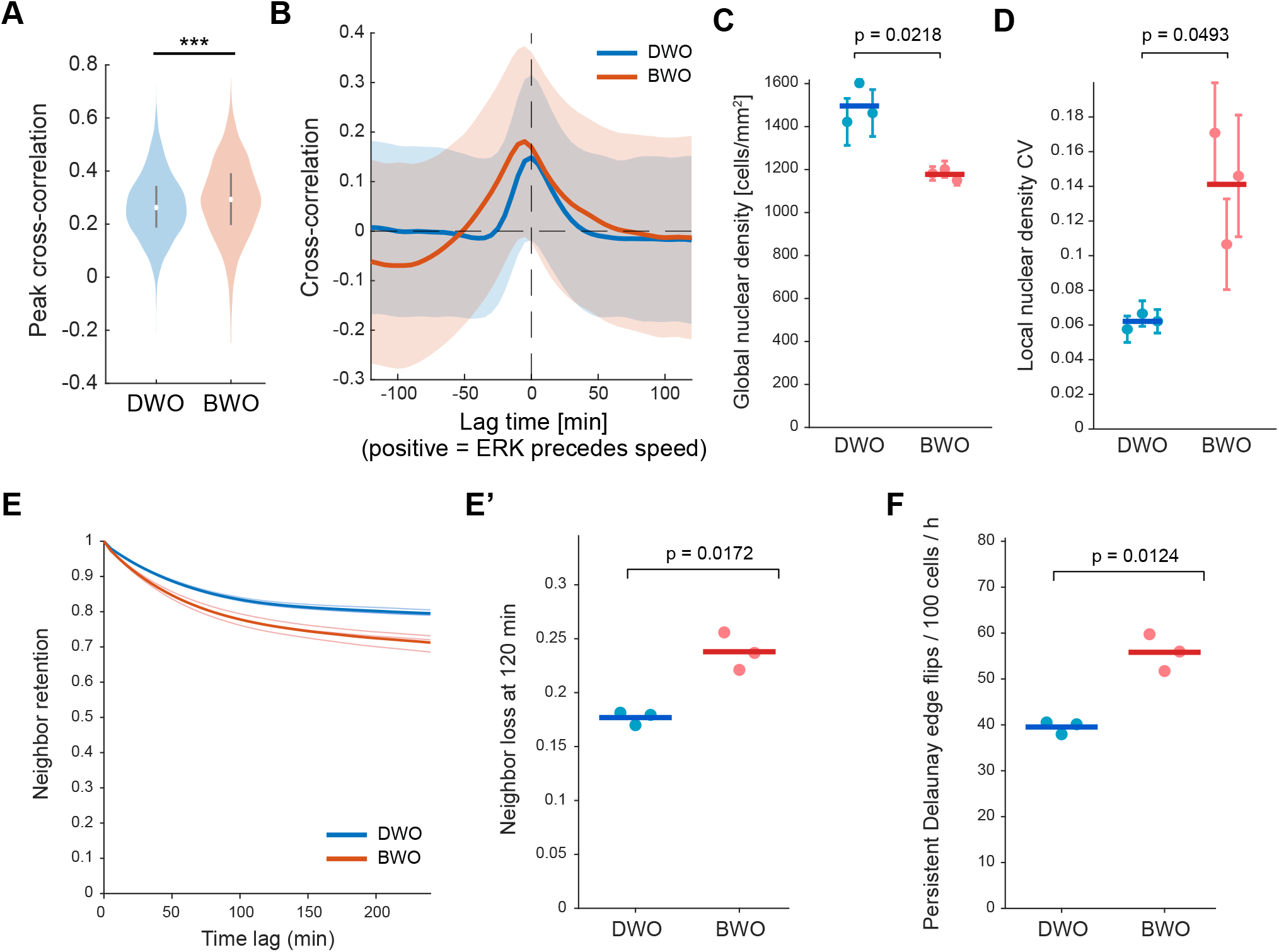
ERK–speed coupling, density heterogeneity, and topological rearrangements. (A) Violin plots showing the distribution of single-cell peak ERK–speed cross-correlation coefficients in DWO and BWO cells. Mean peak cross-correlation values were 0.27 for DWO cells (n = 3,133 cells) and 0.29 for BWO cells (n = 3,066 cells); two-sample t-test, p = 3.16 × 10^-12^. (B) Mean ERK–speed cross-correlation as a function of lag time in DWO and BWO cells. Solid lines indicate the mean across cells, and shaded regions indicate standard deviation. The dashed vertical line indicates zero lag. Positive lag indicates that ERK activity precedes cell speed. (C) Global nuclear density in DWO and BWO monolayers. Each circle represents the temporal mean from one independent experiment over the 480-min observation period; error bars indicate the temporal s.d., and horizontal bars indicate the mean of biological-replicate means. DWO, N=3; BWO, N=3. Welch’s two-sample t-test on biological-replicate means, p=0.0218. (D) Spatial heterogeneity of nuclear density in DWO and BWO monolayers, quantified as the coefficient of variation (CV) of local nuclear density. Each circle represents the temporal mean from one independent experiment; error bars indicate the temporal s.d., and horizontal bars indicate the mean of biological-replicate means. DWO, N=3; BWO, N=3. Welch’s two-sample t-test on biological-replicate means, p=0.0493. (E,E’) Neighbor retention and loss in DWO and BWO monolayers. (E) Neighbor retention as a function of time lag, showing a faster loss of initial neighbors in BWO cells. (E’) Neighbor loss at 120 min. Mean neighbor loss was 0.177±0.006 for DWO and 0.238±0.017 for BWO. N=3 independent experiments; Welch’s two-sample t-test on biological-replicate means, p=0.017. (F) Rate of persistent T1-like Delaunay edge flips. The event rate increased from 39.5 ± 1.4 events per 100 cells per hour in DWO to 55.8 ± 4.0 events per 100 cells per hour in BWO (N=3 independent experiments; Welch’s two-sample t-test, p=0.012). Gray circles indicate individual experimental means; horizontal bars indicate the mean across experiments.

**Figure S5.**
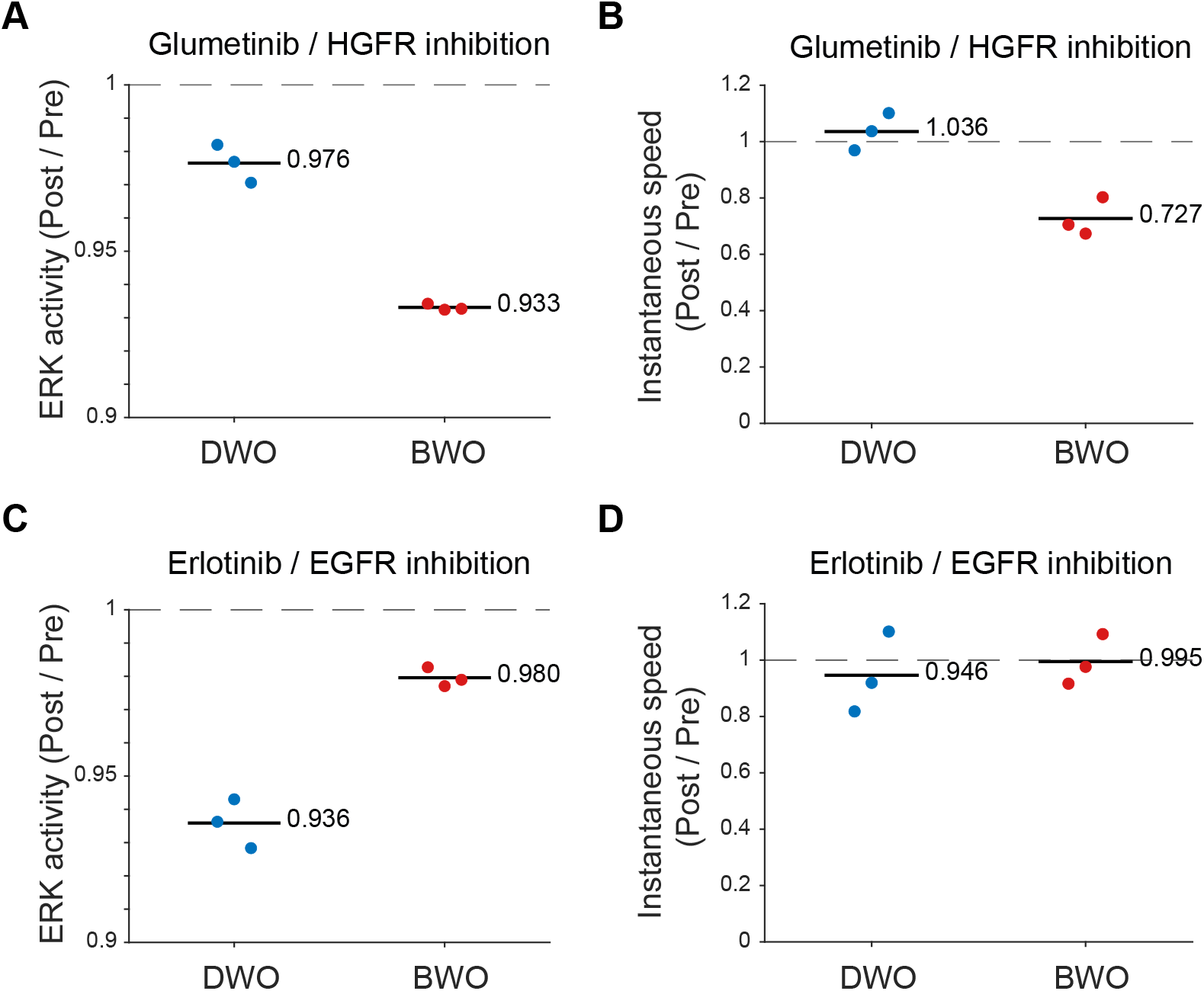
Quantification of responses to HGFR and EGFR inhibition. (A,B) Relative changes in ERK activity (A) and instantaneous cell speed (B) following Glumetinib treatment in DWO and BWO cells. (C,D) Corresponding changes following Erlotinib treatment. Values are expressed as post-/pre-treatment ratios; each point represents one analysis group, horizontal bars indicate the mean, and the dashed line denotes no change (ratio = 1).

**Figure S6.**
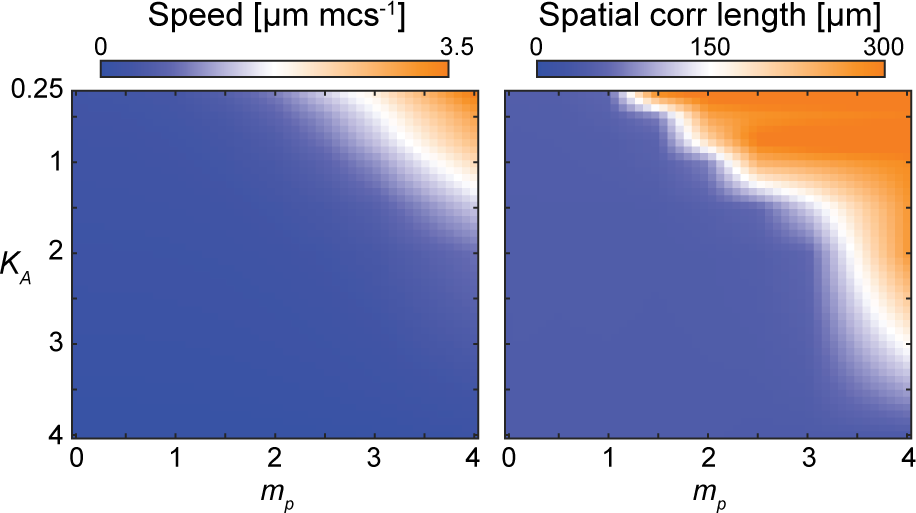
Phase diagrams of collective migration speed and spatial correlation length. Instantaneous cell speed (left) and spatial velocity correlation length (right) as functions of protrusion strength *m_p_* and cell elasticity *K_A_* in the protrusion-driven regime, with contraction strength set to zero. Warmer colors indicate larger values.

## SUPPLEMENTARY VIDEOS

**Video 1** Optical flow in expanding epithelial monolayers of DWO (top) and BWO (bottom) cells. See Figure 3A for the optical flow color wheel. Scale bar, 100 µm.

**Video 2** ERK activity in expanding epithelial monolayers of DWO (top) and BWO (bottom) cells. See Figure 3D for the ERK activity colormap. Scale bar, 100 µm.

**Video 3** Optical flow in confluent epithelial monolayers of DWO (left) and BWO (right) cells. See Figure 4A for the optical flow color wheel. Scale bar, 200 µm.

**Video 4** ERK activity in confluent epithelial monolayers of DWO (left) and BWO (right) cells. See Figure 4A for the ERK activity colormap. Scale bar, 200 µm.

**Video 5** Glumetinib (HGFR inhibitor) treatment in DWO (left) and BWO (right) cells. Time zero indicates treatment onset. Scale bar, 100 µm.

**Video 6** Erlotinib treatment (EGFR inhibitor) in DWO (left) and BWO (right) cells. Time zero indicates treatment onset. Scale bar, 100 µm.

**Video 7** CK-666 (Arp2/3 inhibitor) treatment in DWO (left) and BWO (right) cells. Time zero indicates treatment onset. Scale bar, 100 µm.

**Video 8** CPM simulation of the contraction-based regime, showing cell morphology with front–rear polarity (left), optical flow (middle), and velocity vector fields (right). Parameters: ***m_c_*** = **2**, ***m_p_*** = **0**, ***K_A_*** = **1**, ***ε*** = **0**. **2**. See Figure 6E for the optical flow color wheel.

**Video 9** CPM simulation of the protrusion-based regime, showing cell morphology with front–rear polarity (left), optical flow (middle), and velocity vector fields (right). Parameters: ***m_c_*** = **0**, ***m_p_*** = **2**, ***K_A_*** = **1**, ***ε*** = **0**. **2**. See Figure 6E for the optical flow color wheel.

## Notes

### Competing Interest Statement

The authors have declared no competing interest.

### Summary of Updates

This revised version includes substantial new experiments and analyses addressing reviewer comments. We added characterization of multinucleation and p53 p21 signaling, longitudinal analysis of the BWO associated adhesion phenotype, density matched measurements of collective migration, quantitative ERK speed cross correlation, and direct analysis of cell neighbor loss and persistent T1 like rearrangements. We also added quantitative comparisons of responses to HGFR and EGFR inhibition and expanded the computational analysis. The interpretation of the BWO state has been refined from a leader like state to a protrusion driven migratory state, and the distinction between persistent collective motion and tissue fluidization is now more clearly defined. The computational model, statistical analyses, Methods, Discussion, and code documentation have also been substantially revised.

